# Draper-mediated efferocytosis by *Drosophila* imaginal disc epithelial cells clears cellular debris during regeneration

**DOI:** 10.64898/2026.05.04.722804

**Authors:** Snigdha Amit Mathure, Kaela Maghinang, Rachel Smith-Bolton

## Abstract

**Article summary:** Tissue regeneration requires organized responses to damage, including clearance of cellular debris. Using a genetic ablation system in *Drosophila* wing imaginal discs, we show that most debris is cleared within two days despite the absence of immune cell recruitment, which is restricted by the basement membrane. In the absence of immune cells, debris clearance occurs through Draper-mediated efferocytosis and lysosomal processing by epithelial cells. Disruption of this pathway delays debris removal and impacts regeneration. Residual debris consists of a heterogeneous mix of cellular components, indicating non-selective clearance. Together, our findings identify epithelial cells as key non-professional phagocytes during regeneration.

Regeneration is a coordinated process that restores tissue integrity following damage. Following injury, tissues initiate early responses, including epithelial remodeling and clearance of cellular debris. However, how debris clearance is coordinated with regenerative growth to ensure efficient tissue repair remains poorly understood. To address how early damage responses, particularly debris clearance, are coordinated with regeneration, we used a genetic ablation system in *Drosophila* wing imaginal discs to induce apoptosis in the pouch region. Targeted damage generates cellular debris that localizes to both the apical and basal sides of the epithelium. We show that most cellular debris is cleared within two days after damage, although some debris persists apical to the regenerating epithelium. Notably, immune cells are not recruited to the damaged tissue due to restricted access by an intact basement membrane. Instead, we discovered that debris clearance is mediated by efferocytosis, whereby neighboring hinge epithelial cells activate JNK signaling and engulf debris via lysosomal formation. Reduction of efferocytosis by mutation of the phagocytic receptor *Draper* delays debris removal and increases debris persistence. This impairment has a modest impact on regeneration, as measured by adult wing size. Finally, our data indicate that residual debris consists of a heterogeneous mixture of cellular components, suggesting no preferential targeting by the clearance machinery. Together, our results reveal a previously unappreciated role for epithelial cells as non-professional phagocytes for debris clearance during regeneration.

## Introduction

Regeneration proceeds through multiple tightly regulated stages, including early damage-sensing responses that initiate regeneration, followed by proliferation and growth, and ultimately repatterning and morphogenesis to restore tissue structure and function (reviewed in Poss and Tanaka 2024; Li et al. 2025). The *Drosophila* wing imaginal disc is a well-established model system for studying tissue regeneration and repair. These epithelial structures possess robust regenerative capacity and high genetic tractability, making them extensively characterized over the past several decades (reviewed in Tripathi and Irvine 2022; Worley and Hariharan 2022). Earlier studies primarily relied on physical fragmentation and ex vivo culture of wing discs to investigate regeneration (reviewed in Fox et al. 2020). More recently, the development of genetic ablation systems has enabled precise and controlled induction of tissue damage, providing a powerful approach to study regenerative responses (Smith-Bolton et al. 2009; Bergantiños et al. 2010). These systems involve targeted expression of pro-apoptotic factors, leading to rapid, localized apoptosis within the epithelium and the generation of substantial amounts of apoptotic cell debris (reviewed in Fox et al. 2020). Damaged wing discs activate wound-responsive mechanisms and multiple molecular pathways that are drivers of regeneration. These drivers include reactive oxygen species (ROS) (Santabárbara-Ruiz et al. 2015; Brock et al. 2017; Khan et al. 2017), Wingless (Wg) signaling (Smith-Bolton et al. 2009; Harris et al. 2016), Jun N-terminal kinase (JNK) signaling (Bosch et al. 2005; Bergantiños et al. 2010), p38 MAPK activity (Santabárbara-Ruiz et al. 2015), JAK/STAT signaling (Katsuyama et al. 2015; La Fortezza et al. 2016), Dpp signaling (Herrera et al. 2013), and Yorkie (Yki) activity (Sun and Irvine 2010; Grusche et al. 2011). While apoptosis-induced damage activates these regenerative growth pathways, the extensive cell death necessitates efficient clearance of apoptotic debris from the tissue.

Rapid and efficient clearance of apoptotic cells is a critical process that maintains tissue homeostasis, suppresses inflammatory responses, and regulates immune signaling (reviewed in Elliott and Ravichandran 2010). Failure to remove apoptotic cells in a timely manner can lead to secondary necrosis and the uncontrolled release of harmful intracellular contents (reviewed in Poon et al. 2014). Thus, apoptosis must be tightly coupled with the swift clearance of apoptotic debris. Clearance of dead or dying cells is typically mediated by specialized immune cells called professional phagocytes (reviewed in Melcarne et al. 2019). These include macrophages in mammals and plasmatocytes in *Drosophila*, which serve as the primary cell types responsible for engulfing and removing cellular debris (reviewed in Shklover et al. 2015; Melcarne et al. 2019). In *Drosophila*, plasmatocytes phagocytose caspase-activated cells within tumor microenvironments, such as in malignant eye imaginal disc tumors induced by oncogenic mutations (Hirooka et al. 2025). Furthermore, clearance of apoptotic corpses by plasmatocytes during *Drosophila* embryonic development represents a well-established paradigm of phagocytosis (Weavers et al. 2016).

In addition to professional phagocytes, non-professional phagocytes, including tissue-resident neighboring cells, also play important roles in engulfing apoptotic material through a process known as efferocytosis (reviewed in Shklover et al. 2015; Melcarne et al. 2019). While phagocytosis broadly refers to sensing and internalizing extracellular particles larger than ∼0.5µm, efferocytosis is a specialized form of phagocytosis dedicated to the recognition and clearance of apoptotic cells (reviewed in Uribe-Querol and Rosales 2020; Moon et al. 2023). In *Drosophila*, this function is widely distributed across non-professional phagocytes in multiple tissues. For example, glial cells contribute to central nervous system remodeling during development by clearing apoptotic neurons (Kurant et al. 2008; Etchegaray et al. 2016), while ovarian follicle epithelial cells engulf nurse cells during oogenesis (Meehan et al. 2016; Serizier and McCall 2017). Similarly, glial cells mediate debris clearance during larval axon pruning (Awasaki and Ito 2004), and epidermal cells remove degenerating dendrites (Han et al. 2014). Notably, in *Drosophila* eye imaginal disc epithelial cells, neighboring cells function as non-professional phagocytes and mediate engulfment of oncogenic cells in neoplastic tumor-suppressor mutant tissues, highlighting a broader role for epithelial tissues in cell clearance (Ohsawa et al. 2011). Whether similar non-professional phagocytic mechanisms operate in regenerating wing imaginal discs remains unclear.

In this study, we investigated how apoptotic cell debris is cleared during wing imaginal disc regeneration. We found that most debris generated by tissue ablation was eliminated within two days post-injury, while a small portion of debris persisted. Notably, *Drosophila* immune cell subtypes were not recruited to the damaged epithelium. To understand this lack of immune cell involvement, we examined tissue accessibility and found that the intact basement membrane of the regenerating wing disc serves as a physical barrier, restricting immune cell entry. In the absence of professional phagocyte recruitment, we show that neighboring undamaged epithelial cells in the hinge region of the wing disc activate JNK signaling and act as non-professional phagocytes, engulfing and processing apoptotic debris through lysosome formation. We impaired debris clearance by using a mutation in the phagocytic receptor *draper*, resulting in increased debris persistence and delayed lysosome formation. This defective debris clearance had minimal impact on overall regeneration, as assessed by adult wing size. Finally, we characterized the composition of debris that persisted more than two days after damage and found it to be a heterogeneous mixture of cellular components and organelles. Together, our results reveal that apoptotic debris clearance is independent of immune cells and instead relies on Draper-mediated efferocytosis by surrounding epithelial cells.

## Results

### Genetic ablation system used to induce tissue damage in wing imaginal discs

To understand the early damage response in the wing imaginal disc, we induced targeted tissue damage specifically in the wing pouch, which gives rise to the adult wing blade. To achieve this damage, we used our genetic ablation system (Smith-Bolton et al. 2009), which combines spatial and temporal control of ablation via GAL4-UAS-GAL80^ts^ to induce tissue damage (Figure 1a). Apoptosis was induced in third-instar larvae by expressing the pro-apoptotic transgene *UAS-reaper* in the *rotund-GAL4*-expressing wing pouch cells. Briefly, the larvae were reared at 18°C, where a *tubulin-GAL80^ts^* inhibited apoptosis by restraining GAL4 from binding and activating expression of *UAS-reaper*. On day 7 after egg lay, the temperature was shifted to 30°C to relieve the inhibition by GAL80^ts^, enabling GAL4-induced Reaper to drive cell death in the wing imaginal disc pouch cells for 24 hours. The animals were then shifted back to 18°C to permit regeneration, and the subsequent recovery timepoints (0 to 72 hours post-damage) were used to assess the process of regeneration in the wing discs, including early damage responses such as debris clearance. Adult wing sizes were used as a readout of the regenerative capacity of the larval wing imaginal discs.

**Figure 1.**
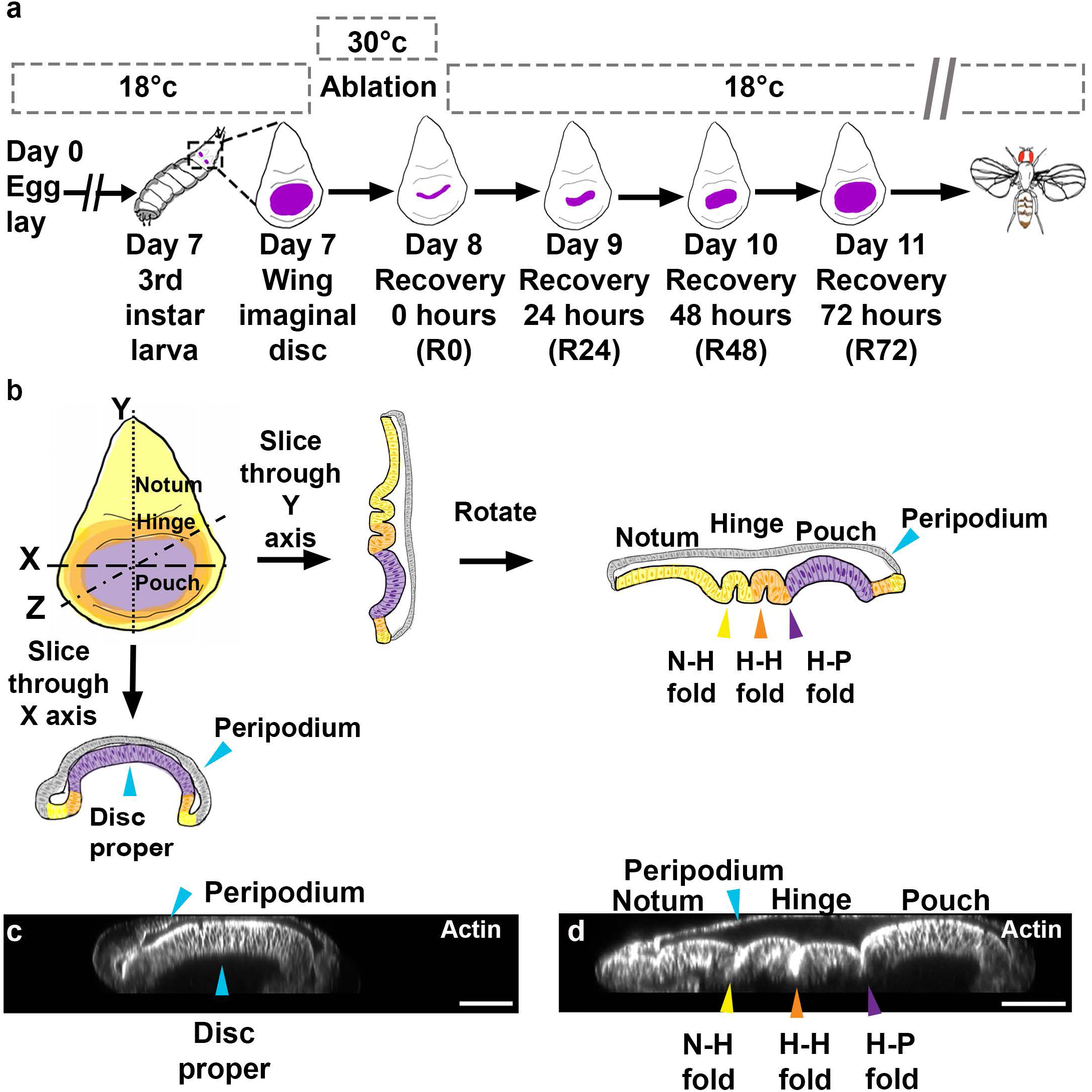
Tissue Damage and Orthogonal Projections of the Wing Imaginal Disc. (a) Schematic representation of the genetic ablation system. (b) Schematic representation of orthogonal projections of wing imaginal disc images, showing morphology in the XZ projection through the pouch and the YZ projection through the notum, hinge, and pouch. Purple cells are the pouch, orange cells are the hinge, yellow cells are the remainder of the disc proper. Yellow arrowhead: Notum-Hinge fold (N-H); Orange arrowhead: Hinge-Hinge fold (H-H); Purple arrowhead: Hinge-Pouch fold (H-P). (c) Phalloidin staining showing actin in the XZ projection of an undamaged wing disc through the pouch. (d) Phalloidin staining showing actin in the YZ projection of an undamaged wing disc through the notum, hinge, and pouch. Yellow arrowhead: Notum-Hinge fold (N-H); Orange arrowhead: Hinge-Hinge fold (H-H); Purple arrowhead: Hinge-Pouch fold (H-P). Scale bars: 50 µm.

To examine early responses to tissue damage, wing imaginal discs were characterized morphologically using confocal microscopy (Figure 1b-d). We used double-sided tape as spacers while mounting to preserve the 3-dimensional architecture of the fixed wing imaginal disc (Aldaz et al. 2010). The epithelial organization of the wing imaginal disc was resolved by orthogonal projection of z-stack images, revealing the squamous peripodium and the columnar disc proper (Figure 1b-d) (reviewed in Tripathi and Irvine 2022). This method of imaging allowed us to observe distinctive folds in the columnar epithelium between the notum, hinge, and wing pouch, denoted as Notum-Hinge (N-H), Hinge-Hinge (H-H), and Hinge-Pouch (H-P) folds (Figure 1b, d).

### Cellular debris is cleared during the first 48 hours of regeneration with some persisting debris

Two of the initial steps in early regeneration are re-establishing tissue continuity and clearing cellular debris (reviewed in Smith-Bolton 2016). To determine how cellular debris is processed, we closely examined the dynamics of debris clearance during wing imaginal disc regeneration. To label cellular debris, we expressed *UAS-EYFP* along with our ablation system, which resulted in EYFP expression in all cells in which GAL4 induced expression of *reaper*. As a result, the EYFP signal was observed in apoptotic cells and the resulting cellular debris. To obtain a comprehensive view of debris clearance, we examined orthogonal slices of confocal image stacks of undamaged and regenerating wing imaginal discs (Figure 2, S1).

**Figure 2.**
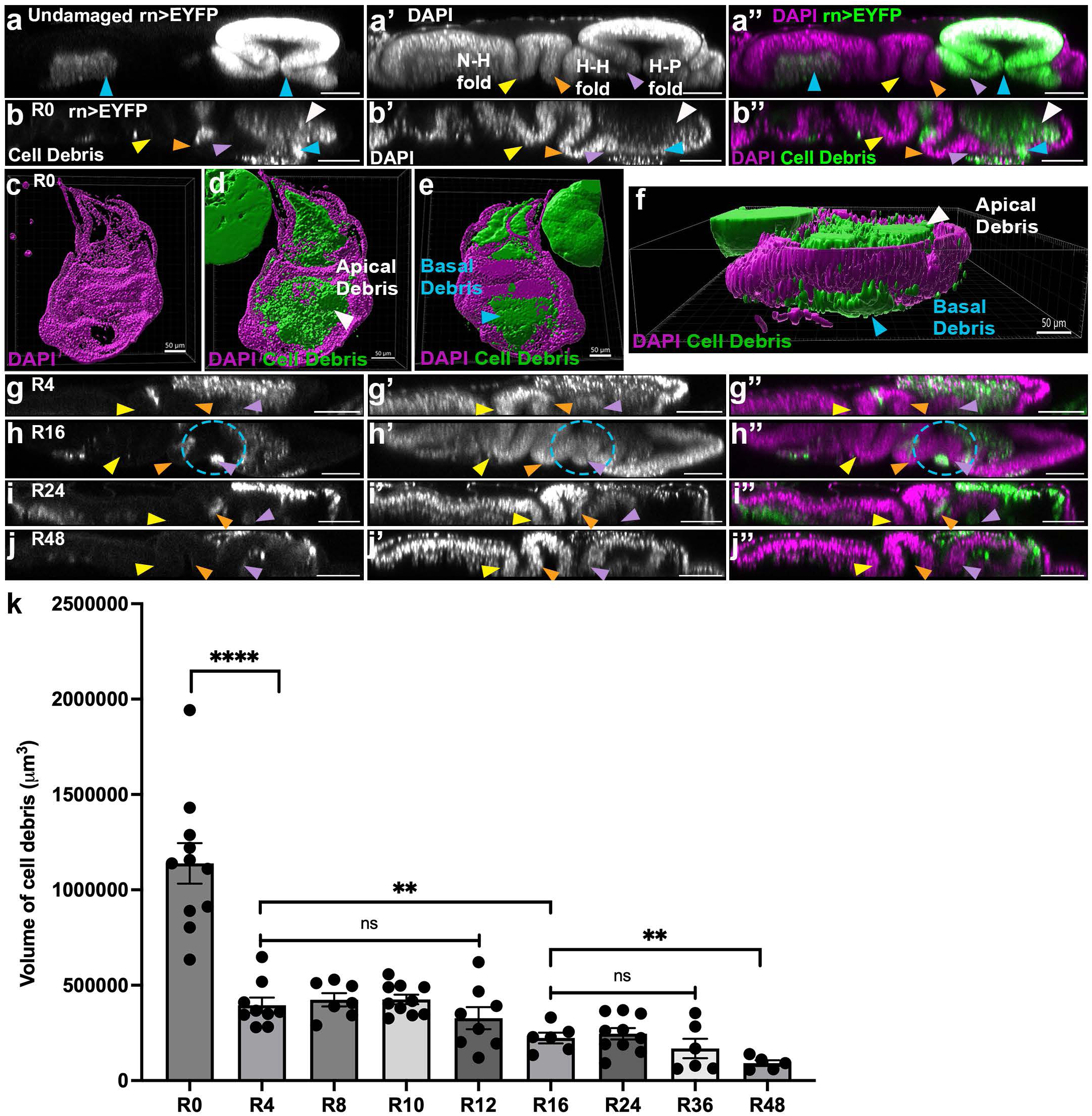
Cellular debris localization and clearance during wing disc regeneration. (a-a″) Orthogonal YZ projection of undamaged wing imaginal disc expressing *rn-GAL4 > UAS-EYFP* (rn>EYFP). *rn*>EYFP expression marks the pouch and adult muscle precursor cells in the notum (blue arrowheads, a and a’’). Nuclei are labeled with DAPI (a′-a″). For all panels, Yellow arrowhead: Notum-Hinge fold (N-H); Orange arrowhead: Hinge-Hinge fold (H-H); Purple arrowhead: Hinge-Pouch fold (H-P). (b-b″) Orthogonal YZ projection of R0 wing disc expressing *rn*>EYFP (b,b’’) with DAPI staining (b′-b″). Apical debris is indicated by white arrowheads, and basal debris by blue arrowheads. (c-f) Three-dimensional reconstructions of R0 wing discs stained with DAPI and expressing *rn*>EYFP to label cell debris. (c-e) Wing disc showing apical debris (white arrowhead) and basal debris (blue arrowhead). (f) Orthogonal side view highlighting the apical and basal localization of debris. (g-g″) Orthogonal YZ projection of R4 wing disc expressing *rn*>EYFP (g,g’’) with DAPI staining (g′-g”). (h-h″) Orthogonal YZ projection of R16 wing disc expressing *rn*>EYFP (h,h’’) with DAPI staining (h′-h”). Debris localized in the hinge epithelium (blue dotted circle). (i-i″) Orthogonal YZ projection of R24 wing disc expressing *rn*>EYFP (i,i’’) with DAPI staining (i′-i”). (j-j″) Orthogonal YZ projection of R48 wing disc expressing *rn*>EYFP (j,j’’) with DAPI staining (j′-j”). (K) Quantification of total cell debris volume across regeneration time points: R0 n = 11 discs, R4 n = 9 discs; ****P = 0.0001 compared to R0, R8 n = 7 discs, R10 n = 5 discs, R12 n = 8 discs, R16 n = 6 discs; **P = 0.0036 compared to R4, R24 n = 10 discs, R36 n = 6 discs, and R48 n = 5 discs; **P = 0.0039 compared to R16. Pairwise comparisons showed no significant differences between R4 and R8 ns, P = 0.6004, R4 and R10 ns, P = 0.5330, R4 and R12 ns, P = 0.3507, R16 and R24 ns, P = 0.5961 or R16 and R36 ns, P = 0.3727. Statistical significance was determined using Welch’s t-test. Error bars: SEM. Scale bars: 50 µm.

In the undamaged control, *rotund*-expressing cells were detected via EYFP signal in the pouch, extending from the H-P fold to the end of the wing pouch (Figure 2a-a”). The EYFP signal was also observed within the adult muscle precursor cells adjacent to the notum, as marked by Cut expression (Figure S1a-a***), a marker for adult muscle precursor cells (Sudarsan et al. 2001). Following damage, at 0 hours of recovery (R0), an extensive amount of cellular debris was detected adjacent to the remaining pouch. The EYFP-labelled cellular debris predominantly localized apically to the epithelium of the disc proper, with a smaller portion extruded basally (Figure 2b-b”). This localization is consistent with our previous observations after ablating most of the wing pouch (Brock et al. 2017), but contrasts with debris localization after ablation of smaller areas of cells, which primarily extrude basally (Bergantiños et al. 2010). Surprisingly, we also observed some debris in the undamaged hinge epithelium marked by the H-H fold (Figure 2b-b’’). 3D renderings of regenerating wing imaginal discs at R0 that excluded the peripodium confirmed this pattern of debris localization (Figure 2c-f).

To measure the amount of cell debris in the regenerating wing pouch over time, we estimated debris volume throughout the regeneration time course by first quantifying the planar area of debris using *UAS-EYFP* signal in the top-down (XY) images and then multiplying it by debris height in orthogonal (YZ) cross-sections. By R4, the amount of debris visible in the orthogonal (YZ) cross-sections was less than that at R0, with some portion of debris still detected within the undamaged hinge epithelium (Figure 2g-g”). We did not detect a statistically significant reduction in the volume of debris from R4 to R12 (Figures S1b-d”, 2k). At R16, we observed a noticeable drop in debris volume, marking a second phase of clearance (Figure 2h-h’’, 2k). By R24, most basal EYFP-labeled debris had been cleared, while the remaining apical debris was confined to the lumen between the peripodium and the disc proper (Figure 2i-i”), consistent with our previous observations at this time point (Brock et al. 2017). The overall debris volume, planar area, and height remained unchanged through R36 (Figure 2k, S1e-g). By R48, most of the cellular debris had been eliminated, although some remnants persisted on the apical side of the columnar epithelium (Figure 2j-j”,k). Taken together, these results suggest that by two days post-damage, cellular debris is largely cleared, although a residual fraction persists in the regenerating wing disc.

### Debris clearance during regeneration occurs independently of immune cell recruitment

We next asked how cell debris is cleared in the regenerating wing imaginal disc. Previous work from our lab showed robust Reactive Oxygen Species (ROS) in both cellular debris and the regenerating epithelium, but no detectable hemocyte recruitment at 24 hours post-damage (R24) (Khan et al. 2017). Because a major portion of cell debris is cleared by R24, we reasoned that if hemocytes contribute to this process, their involvement would occur earlier. Therefore, we examined earlier time points to determine whether any of the three major *Drosophila* hemocyte classes, plasmatocytes, lamellocytes, or crystal cells (reviewed in Mathey-Prevot and Perrimon 1998), are recruited to the damaged epithelium.

To assess plasmatocyte recruitment, we used the plasmatocyte-specific reporter Hemolectin-RFP (Hml-RFP) (Makhijani et al. 2011), and immunostained for the plasmatocyte marker Nimrod (NimC) (Kurucz, Váczi, et al. 2007; Kurucz, Márkus, et al. 2007) (Figure S2). Although a small number of *Hml-*positive plasmatocytes were present in the undamaged control discs, their numbers were considerably reduced at R0 and were not significantly different than the controls at R24 (Figure S2a-d). To assess plasmatocyte presence specifically in the regenerating pouch, we quantified *Hml*-positive cells in the pouch region, marked using the *UAS-EYFP* debris reporter. This analysis revealed no significant differences across conditions (Figure S2e).

Similar results were observed with immunostaining for anti-NimC, where we detected a few NimC-positive cells in the undamaged control discs (Figure S2 f-f’). At R0, the number of NimC-positive cells was significantly reduced and was not significantly different than controls at R24 (Figure S2 f-j), consistent with our previous report (Khan et al. 2017).

A recent report showed that the detection of hemocytes in the wing imaginal disc can be influenced by immunostaining procedures, with hemocytes getting dislodged by rigorous washing (Zhu et al. 2025). Therefore, we repeated the NimC experiments without subjecting the discs to vigorous washes after dissection and after incubation with primary and secondary antibody (Figure 3). Although some NimC-positive plasmatocytes were detected in the no-wash condition at both R0 and R24 in the damaged pouch, these numbers were not significantly different from the respective undamaged controls at 0 and 24 hours after thermal shift (Figure 3a-e). The orthogonal YZ slices showed that NimC-positive plasmatocytes were attached to the basal side of the disc proper in the undamaged discs. In contrast, at R0 and R24, plasmatocytes were predominantly localized to the basal side of the peripodial epithelium, a region that lacks cell debris localization (Figure 3f-h). The overall number of NimC-positive plasmatocytes across the whole disc was not significantly different in the no-wash condition when comparing the R0 and R24 regenerating discs to their respective controls (Figure S3a). Additionally, we observed no significant differences in the number of NimC-positive plasmatocytes in the undamaged pouch between the wash and no-wash conditions (Figure S3b). While a significantly higher number of NimC-positive cells was detected at R0 in the no-wash condition compared to the wash condition (Figure S3b), the number was not significantly different from the undamaged no wash condition. Together, these results show no detectable increase in plasmatocytes in the regenerating wing imaginal disc, while suggesting that the plasmatocytes present at damaged and undamaged discs at R0 may be more easily dislodged.

**Figure 3.**
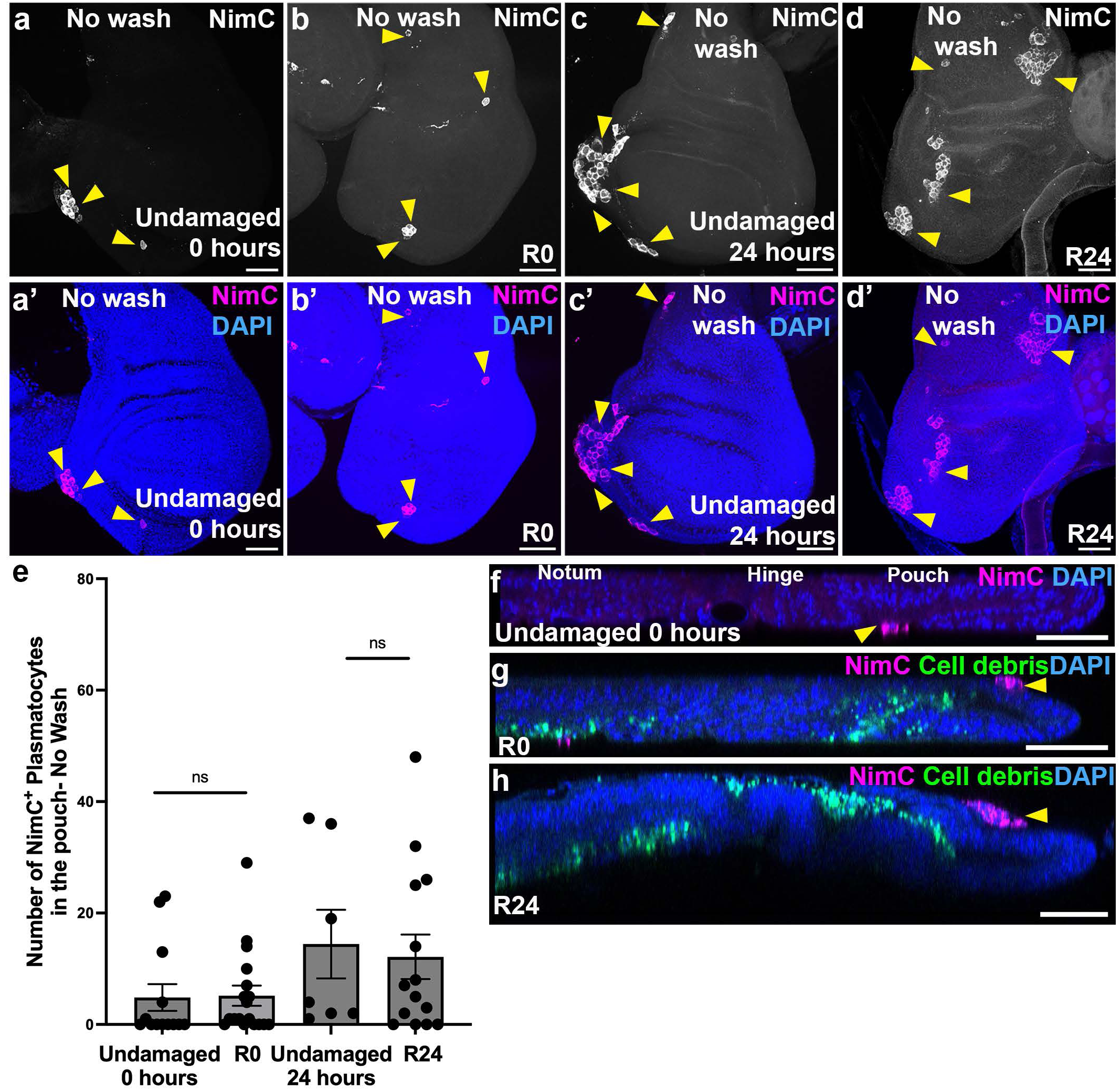
Immune cell recruitment is not detected under no-wash conditions in the regenerating wing disc. (a-d) Anti-Nimrod (NimC) (yellow arrowheads) and DAPI staining under no-wash conditions of an undamaged wing disc at 0 hours (a-a’), a regenerating disc at R0 (b-b′), an undamaged disc at 24 hours (c-c′), and a regenerating disc at R24 (d-d′). (e) Quantification of NimC-positive plasmatocytes within the wing disc pouch under no-wash conditions. Undamaged discs at 0 hours n=13 discs, regenerating discs at R0 n=18 discs, ns, p=0.9158, undamaged discs at 24 hours n=7 discs, and regenerating discs at R24 n=14 discs, ns, p=0.7614. (f-h) Orthogonal YZ projections of discs stained with anti-Nimrod (NimC) to mark plasmatocytes, rn>EYFP to mark cell debris, and DAPI staining to mark nuclei. (f) Undamaged disc at 0 hours. (g) Damaged discs at R0 (h). Damaged discs at R24. Yellow arrowheads indicate NimC-positive plasmatocytes. Statistical significance was determined using Welch’s t-test. Error bars: SEM. Scale bars: 50 µm.

We next sought to detect the presence of lamellocytes using established markers (reviewed in Evans et al. 2014): Atilla/L1 (Kurucz, Váczi, et al. 2007; Evans et al. 2014)*, Atilla-MiET1,* a GFP reporter element insertion which allows in vivo detection of lamellocyte differentiation (Honti et al. 2009), *ItgαPS4* (Irving et al. 2005; Krzemień et al. 2007), and *MSNF9* (Tokusumi et al. 2009). To validate Atilla antibody specificity for lamellocytes, we immunostained third-instar larval hemolymph with Atilla antibody and confirmed that it selectively labels lamellocytes (Figure S4a-a’), which are larger than 40 µm, consistent with previous reports (reviewed in Lan et al. 2020). We examined regenerating and control wing discs and found no Atilla-positive lamellocytes in the undamaged, R0 or R24 discs. However, the antibody marked discs in a consistent speckled pattern within the disc and debris (Figure S4b-d’). To confirm what lamellocytes would look like in the wing imaginal discs, we used a model that overproduces lamellocytes, caused by the gain-of-function allele *hop^[Tum]^* (Luo et al. 1995), which encodes a constitutively active form of the *Hopscotch* (*Hop*)-encoded Janus kinase (JAK). Larvae carrying this mutation develop a melanotic tumor phenotype characterized by extensive lamellocyte differentiation in a temperature-dependent manner and presence of lamellocytes throughout the animal, including at the imaginal discs. We observed lamellocytes in the hemolymph samples as well as on the wing imaginal discs from *hop^[Tum]^* larvae by immunostaining with the Atilla antibody (Figure S4e-f’). No lamellocytes were detected in undamaged or R0 discs under no-wash conditions using the anti-Atilla antibody (Figure S4g-g′). We further examined lamellocyte markers αPS4-GFP and MSNF9mo-GFP, confirming their specificity by co-staining with the Atilla antibody (Figure S5a-b″). Consistent with these results, no lamellocytes were detected in undamaged or R0 discs using αPS4-GFP, MSNF9mo-GFP, or the Atilla-MiET1 reporter (Figure S5c-i). Together, these findings indicate that lamellocytes are not recruited to damaged wing discs for debris clearance.

We then assessed the presence of crystal cells using the crystal cell markers PPO1 (reporter line PPO1-mcherry.F6) (Tokusumi et al. 2017), and Lozenge (Lz) (Lebestky et al. 2000). A few PPO1-mcherry.F6 positive cells were detected in both undamaged control discs and regenerating discs at R0 (Figure S6a-b’). However, their numbers were not significantly different between undamaged and R0 conditions in either the whole disc or the regenerating pouch (Figure S6c-d). By contrast, neither the anti-Lz immunostaining nor the GFP-tagged Lz protein detected any crystal cells in the regenerating discs (Figure S6e-h’). Thus, crystal cells are not significantly increased in number in the regenerating wing imaginal discs.

Finally, to confirm the absence of hemocyte recruitment to the debris region of the regenerating wing imaginal discs, we tested two pan-hemocyte markers: Hemese (He) (Kurucz et al. 2003), and Serpent (Srp) (reviewed in Evans et al. 2014). Immunostaining with an anti-Hemese antibody revealed no significant difference between the number of He-positive hemocytes, both in the pouch and throughout the disc, at R0 and R24 compared to the undamaged controls under the no-wash conditions (Figure S7a-e). To further evaluate hemocyte presence, we verified the pan-hemocyte reporter *srpHemo-H2A::3xmCh*, where the *srpHemo* promoter drives a fusion of 124 amino acids of Histone H2A linked to 3xmCherry, concentrating the fluorescence in the nucleus, and *SrpHemo-3xmCh*, where the *srpHemo* promoter drives a fusion of three copies of mCherry that localizes to the cytoplasm (Gyoergy et al. 2018). These transgenes are reported to detect all three hemocyte sub-classes: plasmatocytes, lamellocytes, and crystal cells. In the third instar larval hemolymph, we found that plasmatocytes positive for NimC (white arrows, Figure S8a-a’, a’’’) and crystal cells positive for Lz-GFP (yellow arrow, Figure S8a’’-a’’’) expressed the *srpHemo-H2A::3XmCh* reporter. Interestingly, we also observed some *srp*-positive cells that were not positive for either NimC1 or Lz (orange arrows, Figure S8a”’). We also detected lamellocytes double-positive for both Atilla and the *srpHemo-H2A::3xmCh* reporter (Figure S8b-b”’). Together, these results indicate that the srpHemo reporters reliably label plasmatocytes, lamellocytes, and crystal cells, while also marking a small population of mCherry-positive cells lacking these three hemocyte markers.

On testing these reporters in regenerating wing discs, we detected *srpHemo-H2A::3XmCh*-positive nuclei at R0, R24, and R48, compared to very few in the undamaged controls (Figure S8c-f). Given that none of the three immune cell types were detected in our previous experiments, we hypothesized that the mCherry-positive nuclei might represent primocytes, a distinct cell type that expresses *srp* but lacks mature hemocyte marker expression (Fu et al. 2020), or they might result from unexpected reporter expression in the ablated cells or the regenerating epithelium. To test whether primocytes were present in the regenerating disc, we stained R48 *srpHemo-H2A::3xmCh* discs with an anti-Antennapedia antibody (Antp), which is a primocyte marker (Fu et al. 2020). We detected no Antp expression in the mCherry-positive nuclei, thereby ruling out the presence of primocytes (Figure S8g-g’).

Orthogonal YZ slices of the srpHemo-H2A::3xmCh discs at R48 revealed mCherry-positive nuclei that colocalized within the regenerating epithelium marked by Discs-large (Dlg) and DAPI (Figure S8h-h’). This unexpected finding prompted further investigation into the identity of the cells containing the mCherry-positive nuclei. Given that hemocytes and columnar epithelial cells are very different in shape, we used a *srpHemo-moe::3xmCh* transgene, in which the mCherry-tagged Moesin would mark the actin cytoskeleton and reveal the morphology of the mCherry-expressing cells (Gyoergy et al. 2018). At R48, *srpHemo-moe::3xmCh* expression showed that these cells had a columnar shape and were integrated into the regenerating epithelium, indicating that they are not immune cells but disc proper cells expressing this reporter (Figure S8i-i’’).

Thus, we are likely observing regeneration-associated activation of the *srp* promoter fragment rather than recruitment or persistence of *srp*-positive hemocytes in the regenerating disc. Collectively, these findings indicate that immune cells are not recruited for debris clearance in the regenerating wing imaginal discs.

### Intact basement membrane restricts immune cell recruitment in the regenerating epithelium

Given our previous analysis showing that immune cells are not recruited to regenerating wing imaginal discs, we asked what factors restrict immune cell recruitment to this tissue. In damaged eye discs, reactive oxygen species (ROS) recruit hemocytes to the site of damage, which then serve as a stimulus to activate JNK signaling in the regenerating epithelium (Fogarty et al. 2016). In regenerating wing discs, although we observe a similar upregulation of ROS and JNK signaling (Khan et al. 2017), we do not detect increased recruitment of immune cells. This discrepancy prompted us to investigate how immune cell recruitment differs between eye and wing discs, despite their overall structural similarity.

A prior study in developing wing discs demonstrated that damage to the basement membrane is sufficient to recruit hemocytes (Diwanji and Bergmann 2020). Using eye imaginal discs, the same group showed that damaged eye discs exhibit basement membrane disruption (Diwanji and Bergmann 2020). To test whether basement membrane integrity similarly regulates immune cell recruitment in regenerating wing discs, we overexpressed matrix metalloproteinase 2 (Mmp2), an endopeptidase that cleaves components of the basement membrane and extracellular matrix (reviewed in Page-McCaw et al. 2007). In undamaged and regenerating wing discs, the basement membrane was visualized using Vkg-GFP, a protein trap including GFP fused to Collagen IV encoded by the *viking* gene (Morin et al. 2001) (Figure 4a-f). At R0, control regenerating discs showed a continuous basement membrane in the pouch region, although Vkg-GFP signal intensity appeared reduced compared to the pouch in undamaged control discs (Figure 4a-b’). In undamaged discs, NimC-positive plasmatocytes were attached to the basal side of either the disc proper or the peripodial epithelium in 11 out of 23 discs (Figure 3f, Figure 4a-a’, d). In the regenerating control discs at R0, NimC-positive plasmatocytes were rarely detected, and, when present, remained associated with the peripodium and did not invade the disc epithelium in 8 out of 24 discs (Figure 3g, Figure 4b-b’,e). By contrast, upon *mmp2* overexpression, we observed disruption of basement membrane at R0, indicated by loss of Vkg-GFP continuity, along with invasion of NimC-positive cells into the disc epithelium (Figure 4c-c’). The presence of immune cell clusters within the damaged pouch in 4 out of 8 discs (Figure 4f) indicates that, in regenerating wing discs, an intact basement membrane functions as a physical barrier to immune cell invasion, distinguishing their damage response from that of damaged eye discs.

**Figure 4.**
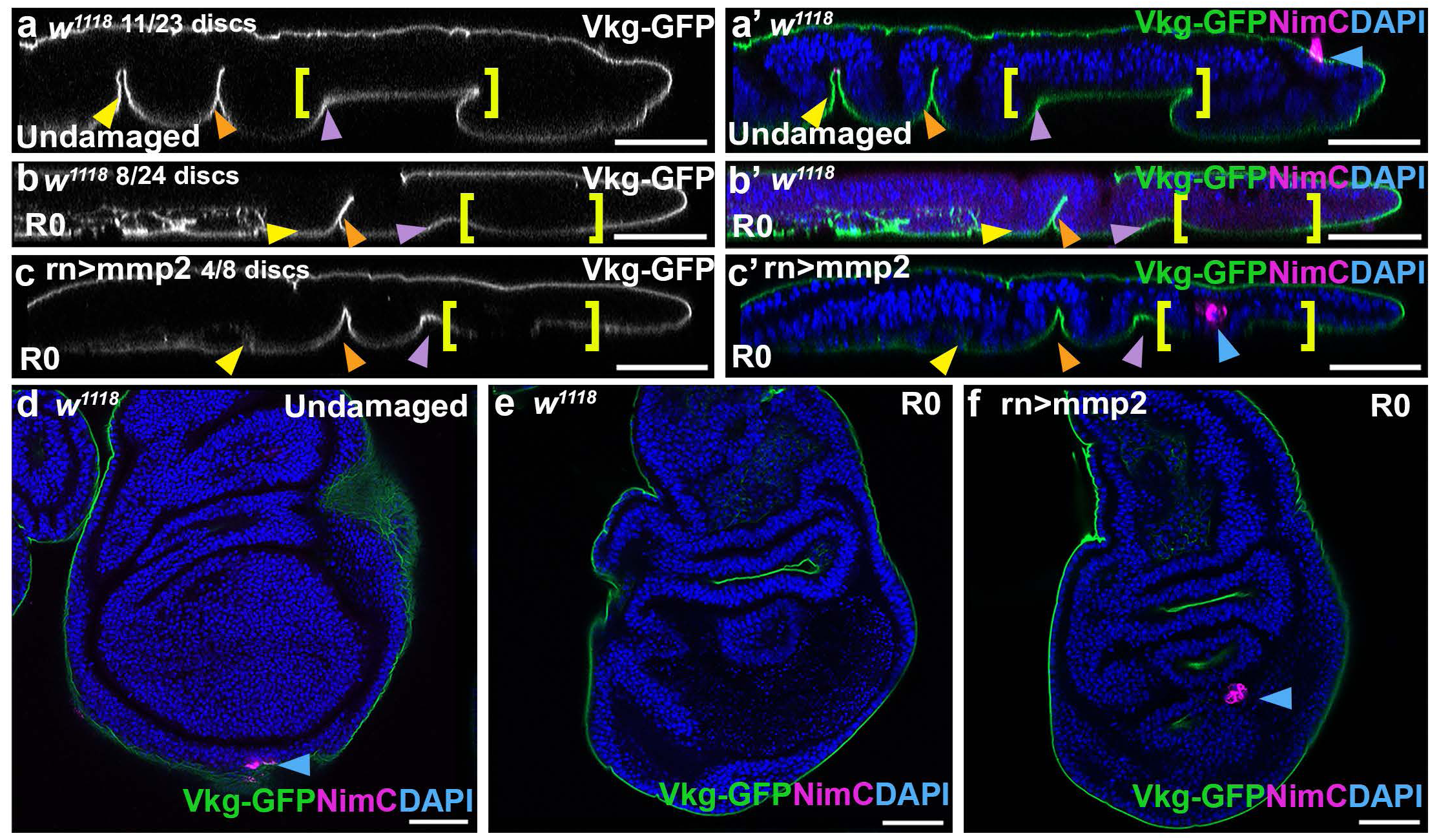
Basement membrane remains intact and restricts immune cell recruitment. (a-c’) Orthogonal YZ projections of wing disc expressing Vkg-GFP and co-stained with NimC and DAPI in control undamaged (*w¹¹¹⁸*) (a-a’), control regenerating R0 disc (b-b’), and *rn*-GAL4 > UAS-mmp2 (*rn*>mmp2) (c-c′) discs. Yellow brackets indicate the wing disc pouch. Blue arrowhead indicates NimC-positive plasmatocytes. For all panels, yellow arrowhead: Notum-Hinge fold (N-H); Orange arrowhead: Hinge-Hinge fold (H-H); Purple arrowhead: Hinge-Pouch fold (H-P). (d-f) Top-down views of undamaged control (d), R0 regenerating disc (e), and *rn*>mmp2 R0 disc (f). Undamaged control n= 23 discs, 11/23 discs were positive for NimC, R0 discs. *w¹¹¹⁸*, n=24 discs, 8/24 discs were positive for NimC, and *rn*>mmp2, n=8 discs, 4/8 discs were positive for NimC. Scale bars: 50 µm.

### Debris clearance during regeneration is associated with lysosome formation in hinge epithelial cells

Having tested the role of hemocytes, the professional phagocytes in *Drosophila*, and finding no evidence for their involvement in debris clearance, we next investigated whether non-professional phagocytes might instead mediate debris clearance during regeneration. Several *Drosophila* cell types can carry out efferocytosis, or phagocytosis of apoptotic debris, including ovarian follicle epithelial cells (reviewed in Serizier and McCall 2017), eye imaginal disc epithelial cells that eliminate oncogenic neighbors (Ohsawa et al. 2011), and embryonic glia (reviewed in Heron et al. 2023). As previously observed, our *UAS-EYFP* labelled debris revealed debris from the damaged pouch within the cells of the adjacent hinge epithelium fold (Figure 5a-a”). This observation is consistent with a role for neighboring hinge epithelial cells in engulfing apoptotic debris generated during pouch ablation. To assess whether there is an increase in phagosomes in the hinge that could be clearing debris, we stained regenerating wing imaginal discs with LysoTracker, a marker of phagosome maturation that labels the completion of the phagocytic engulfment process (Awasaki and Ito 2004; Kurant et al. 2008). We detected very few LysoTracker-positive puncta in both top-down views and orthogonal YZ slices in the undamaged discs (Figure 5b, Figure S9a-a″). Upon damage, LysoTracker-positive puncta were detected in R0 discs, specifically in the hinge-pouch fold (Figure 5c-e’). Quantification showed that damaged R0 discs contained significantly more LysoTracker-positive puncta compared to undamaged controls or R24 discs (Figure 5f-g).

**Figure 5:**
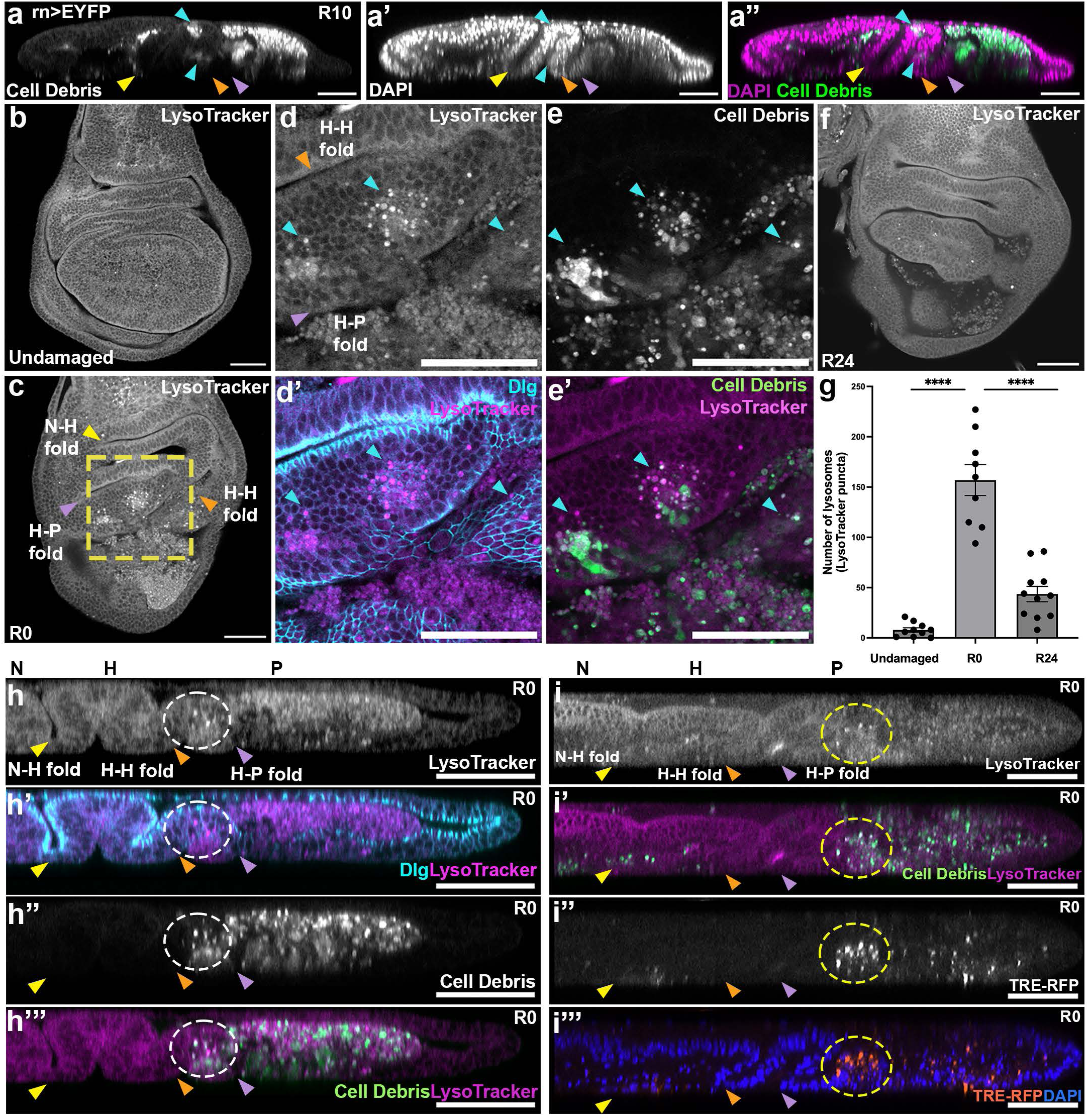
Epithelial debris is cleared through hinge-mediated efferocytosis and lysosome formation. (a-a’’) Orthogonal YZ projection of R10 wing disc expressing *rn*>EYFP with DAPI staining. Blue arrowheads indicate EYFP-labelled cell debris in the hinge epithelium. For all the panels, yellow arrowhead: Notum-Hinge fold (N-H); Orange arrowhead: Hinge-Hinge fold, purple arrowhead: Hinge-Pouch fold (H-P). (b, c) Top-down single-slice views of undamaged control and R0 wing discs stained with LysoTracker. Yellow dashed box-higher magnification region of interest (d-e) Higher-magnification views of the R0 disc in (c) (yellow dashed box) showing LysoTracker-positive puncta (d,d’,e’), anti-Discs large (Dlg) to show epithelial membranes (d’), and *rn*>EYFP-labeled debris (e,e’). Orange arrowhead: Hinge-Hinge fold, blue arrowheads indicate lysosomes. (f) Top-down single-slice view of an R24 wing disc stained with LysoTracker. (g) Quantification of lysosome number (LysoTracker-positive puncta) across regeneration time points. Undamaged control n=10 discs, R0 n=9 discs, ****P= 0.0001 compared to undamaged control, R24 n=11 discs, ****P=0.0001 compared to R0. (h-h‴) YZ orthogonal projection of R0 disc stained with LysoTracker (h,h’,h’’’), Dlg (h’), and expressing *rn*>EYFP to label debris (h’’). The white dotted circle marks lysosomes localized to the hinge-pouch (H-P) fold. (i-i‴) YZ orthogonal projection of R0 disc expressing *rn*>EYFP to mark the debris (i’) and TRE-RFP to mark JNK signaling (i’’,i’’’), and stained with LysoTracker (i,i’) and DAPI (i’’’). The yellow dotted circle marks lysosomes localized in the H-P fold that are associated with JNK signal activation. Error bars: SEM. Scale bars: 50 µm.

Closer examination of R0 discs, with the epithelium marked by Discs large (Dlg), revealed that these lysosomes were confined to the neighboring hinge-pouch epithelial fold (Figure 5d-d’, e-e’). LysoTracker-positive puncta overlapped with the *UAS-EYFP*-labeled debris and were colocalized within the Dlg-marked epithelium as seen by both top-down and YZ orthogonal slices (Figure 5d’-e’, h-h”’).

To refine our understanding of the spatial and temporal dynamics of debris engulfment and lysosome formation during regeneration, we analyzed additional regeneration time points. At R4 and R8, LysoTracker-positive puncta were detected in the hinge epithelium, where they colocalized with the EYFP-labeled debris (Figure S9b-c”). By R10 and R12, puncta persisted primarily within the hinge-hinge and hinge-pouch epithelial folds (Figure S9d-e”). By R24, we observed a significant reduction in the number of LysoTracker-positive puncta relative to R0 (Figure 5e-f, and S9f-f”), consistent with our previous finding that cellular debris volume declines by this stage. By R48, LysoTracker-positive signal was largely absent, as confirmed by orthogonal YZ imaging (Figure S9g-g”).

To confirm the efferocytosis function of the underlying hinge epithelial cells, we tested the genetically encoded fluorescent reporter CharON (Caspase and pH-Activated Reporter, Fluorescence ON), which enables tracking of emerging apoptotic debris and its clearance by efferocytosis (Raymond et al. 2022). Although this reporter has previously been used to detect clearance of apoptotic debris by phagocytes in the *Drosophila* embryo (Raymond et al. 2022), we applied it here to the regenerating wing imaginal discs. The *UAS-CharON* dual reporter consists of a pH-tolerant pH-Caspase-GFP domain that reports apoptosis and a pHlorina red-fluorescent pH sensor that exhibits increased fluorescence upon acidification during apoptotic debris internalization (Raymond et al. 2022). At R0, we detected a pHlorina signal localized to the hinge-pouch fold, as observed in YZ orthogonal sections (10/17 discs), where it colocalized with the pH-Caspase-GFP signal (Figure S9h-h″), consistent with efferocytic processing of apoptotic debris. However, signal intensity was variable when using the CharON reporter.

In eye imaginal discs, activation of JNK signaling in neighboring epithelial cells promotes the elimination of neoplastic tumor cells by engulfment (Ohsawa et al. 2011). To test whether JNK signaling occurs in hinge-pouch epithelial cells during debris clearance in regenerating wing discs, we used the transcriptional JNK signaling reporter *TRE-RFP* (Chatterjee and Bohmann 2012). At R0, we observed localization of both LysoTracker signal and *TRE-RFP* expression within hinge-pouch fold epithelial cells (Figure 5i-i”’). Together, these results support a model in which, after tissue ablation, activation of JNK signaling in neighboring hinge epithelial cells promotes lysosome formation and epithelial engulfment, leading to efficient clearance of cellular debris.

### Reduction of engulfment receptor Draper impairs cell debris clearance and lysosome formation in regenerating wing discs

Next, we asked whether reducing the uptake of cellular debris affects its clearance and subsequent regeneration of the damaged disc. We focused on the phagocytic receptor Draper (Drpr), which is the *Drosophila* ortholog of CED-1, a transmembrane receptor required for engulfment of neurons (Freeman et al. 2003), bacteria (Cuttell et al. 2008), severed axons (MacDonald et al. 2006), germline cells in the ovary (Etchegaray et al. 2012), and imaginal disc cells during cell competition (Li and Baker 2007). Although multiple engulfment receptors have been described in *Drosophila*, we focused on Draper because of its well-established role in mediating engulfment across diverse developmental and injury contexts (Serizier and McCall 2017).

We used the *drpr^Δ5^* null allele(Freeman et al. 2003) to impair Draper function. First, we confirmed that Draper protein levels were reduced in a heterozygous *drpr^Δ5^ /+* background by immunostaining with Draper antibody (Figure S10a-c). We then examined the effects of Draper reduction on debris clearance during wing disc regeneration by quantifying the volume of EYFP-labeled cellular debris at multiple regeneration time points (Figure 6a-g). At R0, regenerating *drpr^Δ5^ /+* discs exhibited a significant increase in debris volume compared to controls (Figure 6a-b’, g). While debris volume decreased over time in control discs, *drpr^Δ5^ /+* discs showed a continued presence of debris at R24, indicating impaired clearance (Figure 6c-d’, g). This defect persisted at R48, when debris was largely eliminated in controls but remained elevated in *drpr^Δ5^ /+* discs, with no significant reduction in volume compared to R0 levels (Figure 6e-f’, g). Consistent with these findings, debris planar area was significantly increased in *drpr^Δ5^ /+* discs compared to controls at all three regeneration time points (Figure S10d). Debris height was comparable between controls and *drpr^Δ5^ /+* mutants at R0; however, at both R24 and R48, debris height was increased in the *drpr^Δ5^ /+* discs, suggesting stalled or delayed debris clearance (Figure S10e).

**Figure 6:**
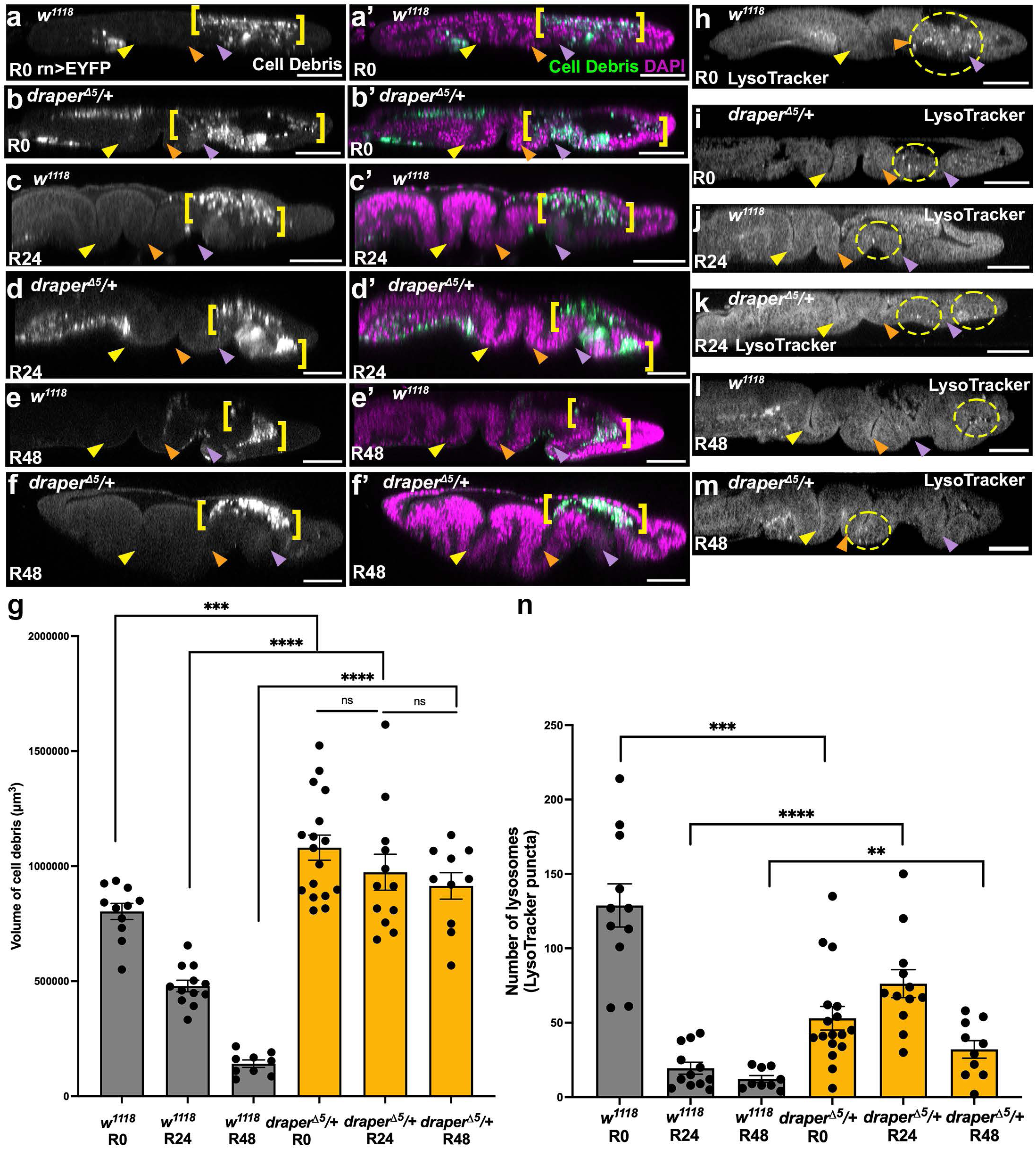
Heterozygous *draper* mutants show impaired debris clearance and altered lysosome dynamics. (a-f) Orthogonal YZ projections of *rn*>EYFP-expressing wing discs stained with DAPI in *w^1118^* controls (a,c,e) and *draper^Δ5^*/+ mutants (b,d,f) at R0 (a-b), R24 (c-d), and R48 (e-f). Yellow brackets indicate *rn*>EYFP-labeled cell debris. For all panels, yellow arrowhead: Notum-Hinge fold (N-H); Orange arrowhead: Hinge-Hinge fold (H-H), and purple arrowhead: Hinge-Pouch fold (H-P). (g) Quantification of debris volume across regeneration time points in *w^1118^* controls (grey bars) and *draper^Δ5^/+* mutants (yellow bars). *w^1118^* R0 n=11 discs, *w^1118^* R24 n=12 discs, *w^1118^* R48 n=9 discs, *draper^Δ5^*/+ R0 n=17 discs, *draper^Δ5^*/+ R24 n=12 discs, *draper^Δ5^*/+ R48 n=10 discs. Pairwise comparisons showed significant differences between control and draperΔ5/+ R0 ***P= 0.0002, R24 ****P= 0.0001, and R48 ****P= 0.0001. No significant differences were found between *draper^Δ5^*/+ R0 and *draper^Δ5^*/+ R24 ns, P=0.2749, and *draper^Δ5^*/+ R24 and *draper^Δ5^*/+ R48 ns, P=0.5491. (h-m) Orthogonal YZ projections of *rn*>EYFP expressing discs, co-stained with LysoTracker and DAPI in *w¹¹¹⁸* controls (h,j,l) and *draper^Δ5^/+* mutants (i,k,m) at R0 (h,i), R24 (j,k), and R48 (l,m). Yellow dotted circles mark lysosome-enriched regions within the hinge and pouch. (n) Quantification of lysosome number (LysoTracker-positive puncta) across regeneration time points in *w^1118^* controls (grey bars) and *draper^Δ5^*/+ mutant wing discs (yellow bars). *w^1118^* R0 n=11 discs, *w^1118^* R24 n=12 discs, *w^1118^* R48 n=9 discs, *draper^Δ5^*/+ R0 n=17 discs, *draper^Δ5^*/+ R24 n=12 discs, *draper^Δ5^*/+ R48 n=10 discs. Pairwise comparisons showed significant differences between the control and *draper^Δ5^*/+ at R0 ***P= 0.0003, R24 ****P= 0.0001, and R48 **P= 0.0090. Error bars: SEM. Scale bars: 50 µm.

We next examined whether impaired debris clearance in *drpr^Δ5^ /+* discs was associated with altered lysosome formation. In control regenerating discs, lysosome number was highest at R0 and progressively decreased at R24 and R48 (Figure 6h-n). By contrast, *drpr^Δ5^ /+* discs exhibited a marked reduction in lysosome number at R0 (Figure 6h-n and S10f-g) followed by an increase at R24 (Figure 6n and S10h-i) and sustained elevated lysosome numbers at R48 compared to controls (Figure 6n, S10j-k). Together, these observations indicate that Draper reduction leads to impaired debris clearance accompanied by a temporal shift in lysosome formation.

To determine whether defective debris clearance impacts regenerative capacity, we assessed pouch regrowth by staining regenerating discs with the wing pouch marker Nubbin (Nub) to quantify pouch area at R48 (Figure S11a-c). We observed no significant difference in Nub-positive pouch area between *drpr^Δ5^ /+* discs and controls (Figure S11c), indicating that growth through this time point is comparable. To understand the effect of impaired debris clearance on regeneration, we quantified adult wing size following disc regeneration (Figure S11d). The adult wings derived from *drpr^Δ5^ /+* revealed a difference in the distribution of adult wing sizes between controls and *drpr^Δ5^ /+* animals, as observed by chi-square analysis. The distribution of adult wing sizes in the mutant showed an increased proportion of severely reduced (0-25%) and reduced (50%) wings (Figure S11d). Taken together, these results demonstrate that epithelial debris clearance during regeneration is at least partially mediated by the phagocytic receptor Draper. Reduction of this Draper-mediated efferocytosis has a modest impact on wing disc regeneration.

### Persistent debris in the regenerating discs consists of heterogeneous cellular components

Our findings thus far indicate that although the majority of cellular debris is cleared by R48 during wing disc regeneration, small remnants of debris persist within the tissue. We therefore sought to determine the composition of this persistent debris. Using *UAS-mCherry* (Ni et al. 2011) to label cell debris, we examined the colocalization of residual debris with markers for different cellular components and organelles. We hypothesized that selective engulfment or clearance might result in preferential persistence of specific cellular constituents.

We did not observe persistence of distinct cellular components within the debris at R48 (Figure 7). Instead, the residual debris consisted of a heterogeneous mixture of cellular materials. Nuclear material (DAPI) (Figure 7 a-a’), nucleolar components (Fibrillarin) (Figure 7 b-b’), nuclear envelope nuclear lamina, as assessed by Lamin A immunostaining (Figure 7c-c’), actin (phalloidin) (Figure 7d-d’), and plasma membrane (CellMask) (Figure 7e-e’) all colocalized with mCherry-labeled debris. Similarly, we examined intracellular organelles, including Mitochondria, Endoplasmic Reticulum (ER), and Golgi, using previously published transgenic EYFP reporters that are under the ubiquitous *spaghetti squash* (*sqh*) promoter (LaJeunesse et al. 2004). Each of the reporters marking mitochondria (Figure 7f-f”), endoplasmic reticulum (Figure g-g’), and Golgi (Figure 7h-h’) colocalized with the mCherry-labelled debris. Collectively, these findings indicate that persistent debris in the regenerating discs does not represent a selectively retained subset of cellular components, but rather a heterogeneous mixture, suggesting no strong preference in the uptake or clearance of specific debris constituents.

**Figure 7:**
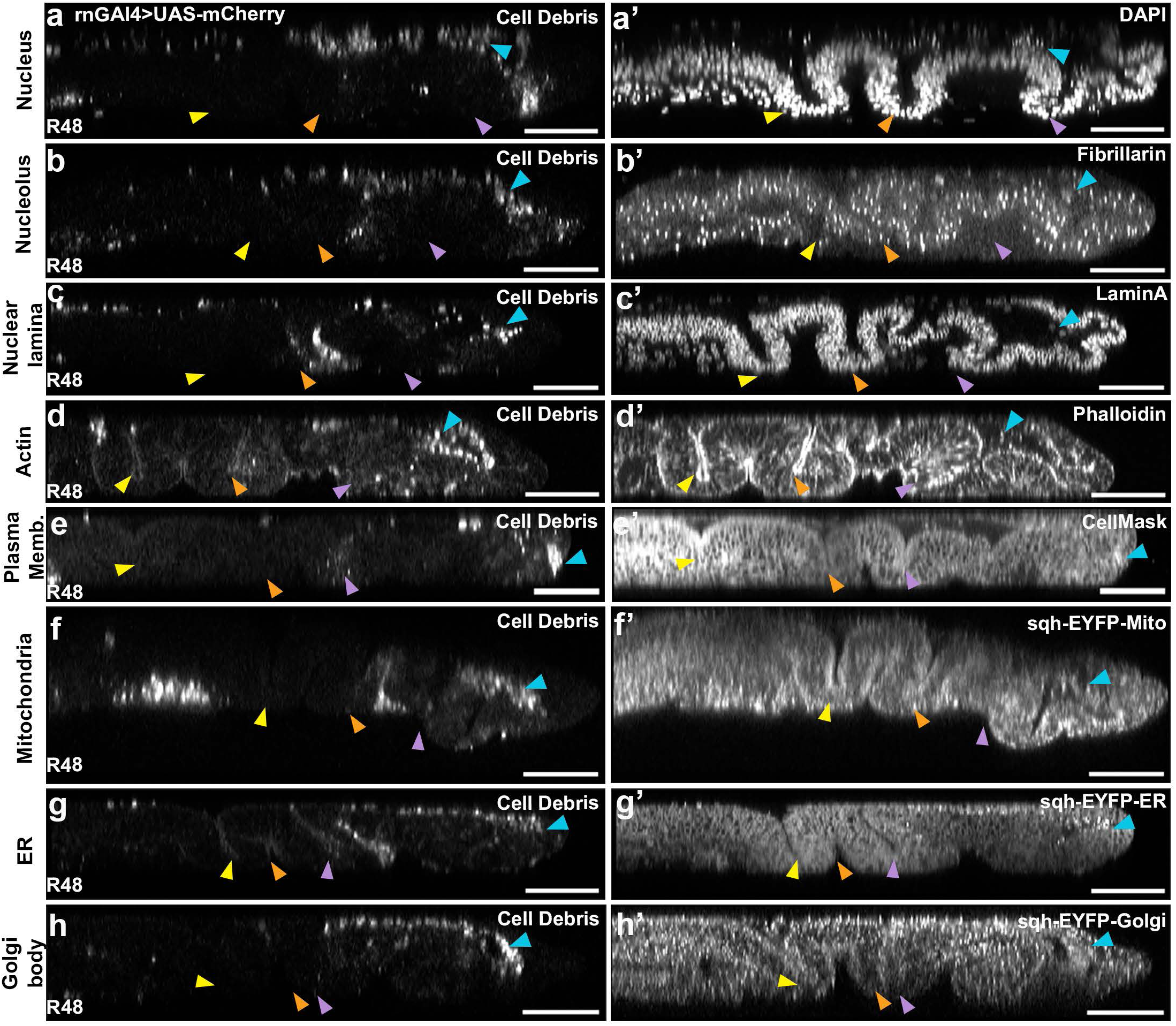
Persistent debris at R48 contains mixed cellular components and organelles. (a-h) Orthogonal YZ projections of R48 wing discs expressing *rn*>mCherry to label cell debris. For all panels, yellow arrowhead: Notum-Hinge fold (N-H); Orange arrowhead: Hinge-Hinge fold (H-H); Purple arrowhead: Hinge-Pouch fold (H-P). (a-a’) Discs were stained with DAPI to mark DNA. 10/10 discs showed DNA in the debris. (b-b’) Discs were stained with Fibrillarin to mark the nucleolus. 3/3 discs showed nucleolus material in the debris. (c-c’) Discs were stained with Lamin A to mark the nuclear lamina. 3/3 discs showed nuclear lamina in the debris. (d-d’) Discs were stained with phalloidin to label F-actin. 3/3 discs showed F-actin in the debris. (e-e’) Discs were stained with CellMask to label the plasma membrane. 3/3 discs showed plasma membrane in the debris. (f-h) R48 discs expressing *rn*>mCherry to label debris were also imaged with transgenic organelle markers. (f-f’) Discs were expressing sqh-EYFP-Mito to label mitochondria. 3/3 discs showed mitochondria in the debris. (g-g’) Discs expressed sqh-EYFP-ER to label the endoplasmic reticulum (ER). 6/6 discs showed ER in the debris. (h-h’) Discs expressed sqh-EYFP-Golgi to label the Golgi apparatus. 10/10 discs showed Golgi in the debris. Blue arrowheads indicate regions of overlap between *rn*>mCherry-labeled cell debris and the indicated cellular components or organelle markers. Scale bars: 50 µm.

## Discussion

In this study, we demonstrate that apoptotic cell debris generated during tissue ablation in the wing imaginal disc is cleared in the absence of immune cell recruitment. Despite robust activation of canonical damage responses, including Wnt/Wg signaling, ROS production, and JNK activation, similar to those observed in damaged eye discs (Fan et al. 2014; Fogarty et al. 2016; Diwanji and Bergmann 2020; Worley and Hariharan 2022), the basement membrane acts as a barrier to immune cell infiltration. In contrast to the eye disc, the regenerating wing disc maintains a continuous extracellular matrix that physically restricts immune cell access to the epithelium. Consistent with this finding, both our study and previous work (Harris et al. 2016) show that the basement membrane remains intact during regeneration. This barrier function is further supported by our prior transcriptional profiling of the regeneration blastema at 24 hours post-damage, which revealed elevated expression of the Collagen IV gene *viking* (log₂FC = 5.45, p = 0.00027) and reduced expression of mmp2 (log₂FC = −1.30, p = 0.0002) (Khan et al. 2017). Together, these transcriptional changes favor basement membrane stabilization over degradation. Importantly, experimental disruption of basement membrane continuity was sufficient to permit immune cell infiltration, directly linking extracellular matrix integrity to immune cell access during epithelial regeneration.

We further show that most apoptotic cell debris generated during tissue ablation is efficiently cleared from the wing imaginal disc within two days after damage by neighboring hinge epithelial cells. This process represents a form of non-professional efferocytosis. Efficient clearance of apoptotic debris is critical for maintaining tissue homeostasis and enabling repair, a process well characterized in mammalian systems (reviewed in Mehrotra and Ravichandran 2022) and during *Drosophila melanogaster* embryogenesis and metamorphosis (reviewed in Adell et al. 2025). When immune cell access is restricted, this function must be carried out by non-professional phagocytes. Thus, the spatial confinement of apoptotic cells within an intact epithelial and basement membrane architecture during wing disc regeneration likely necessitates the uptake of debris by neighboring epithelial cells. Our findings establish the regenerating wing imaginal disc as a model for epithelial efferocytosis, enabling future study of the mechanisms involved and how some debris is left behind. While JNK signaling is a key driver of regenerative responses (reviewed in Tripathi and Irvine 2022), our data indicate that its activation is not confined to the regeneration blastema but also occurs in surrounding epithelial cells. Notably, our results show that JNK activity is elevated in undamaged hinge cells at early time points (R0), whereas at later stages (R24), it is enriched in the regenerating blastema (Brock et al. 2017; Khan et al. 2017), suggesting temporally and spatially distinct activation across undamaged and regenerating cell populations. This early activation in hinge cells suggests that JNK may promote efferocytosis in neighboring epithelial cells, analogous to its role in tumor cell engulfment in eye imaginal discs (Ohsawa et al. 2011).

Delayed or defective clearance of apoptotic debris is associated with secondary necrosis and can lead to pathological outcomes, including autoimmune and chronic inflammatory disorders (reviewed in Kawano and Nagata 2018; Doran et al. 2019). *Draper*, the *Drosophila* homolog of Ced-1, is a well-established receptor required for the recognition and clearance of apoptotic corpses and plays critical roles in multiple developmental contexts (reviewed in Melcarne et al. 2019). In regenerating wing discs, blocking debris clearance with *draper* mutants led to delayed removal of debris and a delayed lysosome formation. Despite these defects, prolonged debris persistence in the regenerating epithelium had only a limited effect on overall regeneration, as assessed by adult wing size. These findings suggest that debris clearance, while actively engaged, may be dispensable during regeneration. However, because *draper* and the ablation system are located on the same chromosome, these experiments were performed in a heterozygous background, and the lack of an effect on adult wing sizes may reflect the incomplete loss of Draper function. Alternatively, compensatory mechanisms involving other engulfment receptors, such as Croquemort, NimC4/SIMU, and integrins (αPS3βPS/βν), which can also mediate apoptotic cell clearance (reviewed in Serizier and McCall 2017; Melcarne et al. 2019), may partially compensate for the reduction in Draper during regeneration. In the regenerating wing disc, persistence of apoptotic debris two days after damage had minimal impact on regeneration of the disc and resulting adult wing sizes.

Residual debris comprises a heterogeneous mixture of cellular components, with no clear bias toward retaining or preferentially clearing specific subcellular structures. In most systems, efferocytic receptors recognize broadly conserved “eat-me” signals on apoptotic material, which may be membrane-anchored or presented via soluble bridging molecules (reviewed in Ravichandran 2010). Phosphatidylserine (PS) is the best characterized of these signals and, when externalized on apoptotic cells, promotes their recognition and clearance (reviewed in Moon et al. 2023). In *Drosophila,* Draper similarly responds to damage-associated cues and PS exposure, enabling the uptake of diverse apoptotic fragments (Tung et al. 2013). Notably, these mechanisms are generally described in terms of broad recognition rather than selective targeting of specific subcellular components, and there is limited evidence supporting strict discrimination between organelle types. Our observations supports a model in which debris clearance is largely non-selective and may be driven by general engulfment mechanisms rather than targeted degradation pathways.

Together, our work redefines debris clearance during regeneration, demonstrating that epithelial cells can autonomously carry out efferocytosis in the absence of immune cell involvement. The wing imaginal disc has not traditionally been considered a model for studying efferocytosis. Our findings now establish apoptosis-induced damage and regeneration in this system as a tractable context to investigate apoptotic debris clearance and epithelial efferocytosis in vivo.

## Materials and methods

### Tissue ablation

All experiments were performed using a protocol adapted from prior work (Smith-Bolton et al. 2009). Egg lays were conducted at 25°C for 4 hours in the dark on grape juice agar plates supplemented with yeast paste (defined as day 0), after which embryos were incubated at 18°C. First instar larvae were collected on day 2 and transferred to Nutri-Fly Bloomington food vials supplemented with yeast paste at a density of approximately 50 larvae per vial. To induce tissue damage, vials were transferred early on day 7 to a water bath maintained at 30°C for 24 hours. Following heat treatment, vials were briefly cooled in an ice-water bath for 1 minute and then returned to 18°C. Regenerating animals were dissected at the indicated time points or allowed to develop to adulthood for quantification of adult wing size. Undamaged control animals were maintained at 18°C and dissected at the third instar crawling stage. In addition, temperature-shifted genetic controls were subjected to the same 30°C heat treatment, but do not undergo ablation.

### Drosophila strains

The fly stocks used for this study are as follows: *w^1118^* (isogenic line made in the Smith-Bolton lab), *w^1118^; rnGal4, UAS-rpr, tubGal80^ts^/TM6B, tubGal80* (Smith-Bolton et al. 2009), *UAS-2xEYFP* (Halfon et al. 2002) (RRID:BDSC_6659), *Hml*Δ*RFP* (gift from K. Bruckner) (Makhijani et al. 2011), *hop^Tum^* (Luo et al. 1995) (RRID:BDSC_8492), *αPS4-GFP* (Sarov et al. 2016) (VDRC_318086), *MSNF9-GFP* (gift from S. Govind) (Tokusumi et al. 2009), *Atilla-MiET1 GFP* (Honti et al. 2009) (RRID:BDSC_23540), *PPO1-mCherry.F6* (Tokusumi et al. 2017) (RRID:BDSC_600219), *lz-GFP* (Kudron et al. 2018), (RRID:BDSC_43954), *srpHemo-3XmCherry/CyO* (RRID:BDSC_78358), *srpHemo-3XmCherry/TM3* (RRID:BDSC_78359), *srpHemo-H2A.3XmCherry/TM3* (RRID:BDSC_78360), *srpHemo-H2A.3XmCherry/CyO* (RRID:BDSC_78361)*, srpHemo-Moe.3XmCherry* (RRID:BDSC_78362) *, srpHemo-Moe.3XmCherry/CyO* (RRID:BDSC_78363) (Gyoergy et al. 2018), *vkg-GFP* (Kyoto DGRC# 110626) (Morin et al. 2001), *UAS-Mmp2* (RRID:BDSC_58706) (Page-McCaw et al. 2003), *TRE-RFP* (gift from D. Bohmann) (Chatterjee and Bohmann 2012), *UAS-CHARON* (gift from W. Wood) (Raymond et al. 2022), *Df(3L)drpr[Delta5]/TM6B* (Freeman et al. 2003), (RRID:BDSC_67033), *sqh-EYFP-Mito* (RRID:BDSC_7194), *sqh-EYFP-ER* (RRID:BDSC_7195), *sqh-EYFP-Golgi* (RRID:BDSC_7193) (LaJeunesse et al. 2004), *UAS-mCherry* (Ni et al. 2011) (RRID:BDSC_35787).

### Immunofluorescence

#### Primary antibodies

Primary antibodies used were mouse anti-cut (1:150, DSHB Cat# 2b10, RRID: AB_528186), mouse anti-Nimrod (1:250, gift from I. Ando) (Kurucz, Márkus, et al. 2007), mouse anti-AtillaL1abc (1:100, gift from I. Ando) (Kurucz, Váczi, et al. 2007), mouse anti-Lozenge (1:10, DSHB, RRID: AB_528346) (Lebestky et al. 2000), mouse anti-Hemese (1:100) (gift from I. Ando) (Kurucz et al. 2003), mouse anti-Antennapedia (1:10, DSHB Cat# 8C11, RRID: AB_528083), mouse anti-Discs large (1:100, DSHB Cat# 4F3, RRID: AB_528203), mouse anti-Nubbin (1:100, DSHB Cat# 2D4, RRID:AB_2722119), rabbit anti-Draper (1:500) (gift from M. Freeman) (Freeman et al. 2003), mouse anti-Fibrillarin (1:100; Abcam, ab4566), mouse anti-Lamin (1:100, DSHB Cat# ADL67.10, RRID: AB_528336).

#### Secondary antibodies

Secondary antibodies used were Alexa Fluor 488, 555, 633, 647 (1:500; Invitrogen, A21424, A21245, A21240, A21071, A21052 and A21094).

#### Stains/Dyes

Phalloidin (1:100, Invitrogen, Thermo Fisher Scientific, R415), DAPI (1:500, Thermo Fisher Scientific, D1306), LysoTracker™ Deep Red (1:250, Invitrogen, Thermo Fisher Scientific, L12492), CellMask™ Deep Red (1:500, Invitrogen, Thermo Fisher Scientific, C10046).

### Immunostaining

Immunostaining was performed using a protocol similar to that previously described (Smith-Bolton et al. 2009) Briefly, dissected larval carcasses were fixed in 4% paraformaldehyde (PFA) in 1× phosphate-buffered saline (PBS) for 20 minutes at room temperature in 1.5 mL microcentrifuge tubes (paraformaldehyde, 16% w/v aqueous solution, methanol-free; Thermo Fisher Scientific, Cat# 043368.9M). Samples were then washed three times for 10 minutes each in 0.1% Triton X-100 in PBS (0.1% PBST) on a nutator at room temperature (Triton X-100; Thermo Fisher Scientific, Cat# PRH5141). Larvae were incubated overnight at 4 °C in primary antibodies diluted as indicated in 0.1% PBST containing 5% normal goat serum (MP Biomedicals™, Cat# MP92939154), followed by three 10-minute washes in 0.1% PBST. Secondary antibodies were used at a 1:500 dilution and incubated overnight at 4 °C. After secondary antibody incubation, samples were washed three times in 0.1% PBST and equilibrated in 70% glycerol in 1× PBS (glycerol, molecular biology grade; Fisher BioReagents™, Cat# BP229-1).

The no-wash immunostaining protocol was adapted from Zhu et al, (Zhu et al. 2025) with a few additions. Following dissections, larval carcasses were fixed in 10 µL of 4% paraformaldehyde (PFA) at room temperature in a 72-well microwell plate (Nunc™ MicroWell™ MiniTrays; Thermo Fisher Scientific, Cat# 438733), with three larval carcasses placed per well. After a 20-minute fixation, carcasses were removed using forceps, briefly dabbed on filter paper (Whatman™ 1001-110 Filter Circles, Cat# 1001-110) to remove excess solution, and transferred to a new well containing 10 µL of 0.1% PBST.

Carcasses were subsequently transferred between primary (1:50) and secondary antibody solutions using repeated filter-paper dabbing in place of conventional wash steps. Primary and secondary antibody incubations were performed overnight, with intermediate 10-minute incubations in 0.1% PBST between antibody steps. Finally, larvae were equilibrated overnight in 70% glycerol in the microwell plate and mounted immediately in VECTASHIELD mounting medium to prevent desiccation due to the small incubation volumes.

### Wing discs mounting

Wing imaginal discs were then dissected and mounted on glass slides (plain microscope slides, Fisher Scientific, Cat# 12550A3; coverslips, 24 × 50 mm, Fisher Scientific, Cat# 12541042) using VECTASHIELD anti-fade mounting medium (Vector Laboratories, Cat# H-1000), and the edges were sealed with Sally Hansen® Quick Dry clear nail polish.

To obtain orthogonal optical sections, a double-sided tape mounting method was adapted from Aldaz et al. (Aldaz et al. 2010) with modifications for fixed-wing imaginal discs. In this modified approach, double-sided tape was used as a spacer. Briefly, double-sided tape (3M 667 Scotch® Double-Sided Tape, Cat# 34-8724-5241-1) was cut into four 19 mm × 2 mm rectangular strips and affixed to a glass microscope slide to make a double-sided tape square. Wing imaginal discs were dissected and mounted at the center of a square coverslip to position the sample as close to the coverslip as possible during imaging (Corning® 18 × 18 mm square #1 cover glass, Cat# 2865-18) in 2 µL of VECTASHIELD mounting medium. The coverslip was then carefully inverted and placed onto the tape-affixed slide. The edges were sealed with Sally Hansen® Quick Dry nail polish to secure the mount and prevent desiccation of the sample.

### Larval hemocytes extraction and immunostaining

Hemocytes were extracted from third instar larval hemolymph using a protocol adapted from Hiroyasu et al. (Hiroyasu et al. 2018). Briefly, a single third instar larva of the indicated genotype was placed in a 2 µL drop of 1× PBS on a circular coverslip (Fisherbrand™ circular cover glass, 18 mm diameter, Cat# 12541005CA) within a glass dissection dish. Larvae were allowed to bleed into the PBS drop for approximately 5 minutes, after which the larval carcass was removed. The hemolymph droplet was then allowed to air-dry on the coverslip for 5 minutes. Coverslips were transferred to a 12-well plate (CytoOne® 12-well plates, Cat# CC7682-7512) and fixed by completely covering the coverslip with 4% paraformaldehyde. Samples were washed three times with PBST on a nutator at room temperature. Primary and secondary antibody incubations were performed overnight at 4 °C on a nutator. Following antibody incubation, coverslips were washed three times for 10 minutes each in PBST, mounted on glass slides with VECTASHIELD mounting medium, and sealed with nail polish.

### Confocal microscopy, image acquisition, and processing

Wing imaginal discs were imaged using a Zeiss LSM 880 or LSM 900 confocal microscope. All images within a given experiment were acquired on the same microscope to ensure consistency using a 20x/0.45 objective. For orthogonal YZ and XZ imaging, z-stacks were collected with a step size of 2 µm. Image acquisition speed was maintained between 3 and 5, and all other imaging parameters, including laser power and detector gain, were kept constant across experimental and control samples.

Orthogonal projections were generated using the Zeiss ZEN Blue software by selecting the ortho view for representative slices. Image files were subsequently opened in ImageJ/FIJI (Schindelin et al. 2012; Schneider et al. 2012) where maximum-intensity projections (MIPs) were generated for downstream processing and figure preparation. 3D reconstruction and surface rendering were performed using Imaris software (Bitplane AG, Oxford Instruments, Zurich, Switzerland), excluding the peripodial epithelium from the z stacks. For Nubbin pouch area quantifications, MIPs were generated, and the Nubbin-positive pouch was manually outlined using the Freehand Selection tool in ImageJ. The enclosed area was then measured and recorded for analysis.

### Cell debris volume quantification

To quantify the volume of cell debris labeled by UAS-EYFP, debris area and height were measured from confocal image stacks. MIPs were generated from confocal z-stacks using ImageJ/Fiji. Top-down XY projections were used to outline debris regions in the UAS-EYFP (green) channel, and the debris area was quantified using the Measure function. Debris height was determined from the same image stacks by generating orthogonal YZ re-slices followed by maximum-intensity projection. Five independent height measurements were obtained per sample using the Straight-Line tool in ImageJ, and the average height within the debris region, as defined by the UAS-EYFP signal, was used as the debris height value. At later regeneration time points, when epithelial regrowth altered tissue morphology, height was measured from YZ projections, accounting for both apical and basal debris relative to the epithelium. Debris volume was estimated by multiplying the measured debris area by the corresponding average debris height for each sample.

### Hemocyte quantification

For quantification of hemocyte number, maximum intensity projections (MIPs) of top-down XY confocal images were generated. Hemocytes were manually counted either across the entire wing disc (including notum, hinge, and pouch regions) or specifically within the rn>EYFP-labeled cell debris in the pouch. For pouch-specific quantification, the rn>EYFP-labelled cell debris was segmented, and immune cells within this domain were counted separately.

### Adult wing mounting and quantification

Adult wings were scored into approximate size categories of 0%, 25%, 50%, 75%, and 100% relative to a wild-type wing that did not undergo damage and regeneration as a disc. Representative wings from each category were dissected and mounted on glass slides using Gary’s Magic Mount. Images were obtained on an Echo Revolve R4 microscope and processed using ImageJ/FIJI. The mounting medium was prepared by dissolving Canada balsam in methyl salicylate at a 5:4 ratio, then gently evaporating the mixture on a medium-heat hot plate with slow stirring at room temperature (for up to 4 days) until most of the solvent had evaporated, leaving a viscous solution. The mounting medium was protected from light during preparation, storage, and mounting.

### Lysosome puncta quantification

Lysosome puncta were quantified using an image analysis pipeline adapted from a previous image analysis pipeline (Gamarra et al. 2020) developed for translation site-specific foci detection and modified here for lysosome analysis. All image processing and quantification were performed using ImageJ/Fiji. Briefly, Lysotracker confocal image stacks were opened in ImageJ/Fiji and converted to grayscale. A region of interest (ROI) encompassing the wing pouch and hinge was manually selected using the freehand selection tool and duplicated, ensuring that all associated hyperstacks were retained. Brightness and contrast were adjusted uniformly across samples to enhance visualization of Lysotracker-positive puncta. A maximum-intensity projection (MIP) was generated from all z-stacks. The MIP image was processed using Process - Filters - Convolve with a 5 x 5 kernel, with the central kernel value set to 48. A threshold was applied to the processed (Image - Adjust - Threshold). Lysosome puncta were quantified using Analyze Particles, with particle size restricted to 2-15 µm² and circularity set to 1.0. Particle counts were exported and plotted using GraphPad Prism.

### Statistical analysis

All statistical analyses were performed using GraphPad Prism. Comparisons between two groups were conducted using Welch’s t-test. Adult wing size distributions were analyzed using a chi-square test. All graphs and plots were generated using GraphPad Prism. All error bars represent the standard error of the mean (SEM).

## Acknowledgments

We thank Dr. Anish Bose, Dr. Felicity Hsu, Connor Powers, Nicholas Magdadaro, Devan Bianchini, Akash Patel, and Shrunali Amin for critical reading of the manuscript. We thank Dr. Katja Brückner, Dr. Shubha Govind, Dr. Dirk Bohmann, Dr. Will Wood, Dr. Marc Freeman, and Dr. Istvan Andó for providing reagents. We acknowledge the Bloomington Drosophila Stock Center (NIH P40OD018537), the Developmental Studies Hybridoma Bank (NICHD, The University of Iowa), the Kyoto Stock Center, and FlyBase for fly stocks, reagents, and database resources. We also thank Dr. Glenn Fried, Dr. Austin Cybersmith, Dr. Kingsley Boateng, Dr. Duncan Nall, and Dr. Umnia Doha of the Core Facilities at the Carl R. Woese Institute for Genomic Biology for assistance with microscopy and image analysis.

## Data and Resource availability

Raw data will be posted to the Illinois Databank (a URL will be generated upon acceptance). All other relevant resources and data can be found within this article and its supplementary information.

---

**Supplementary Figure S1.**
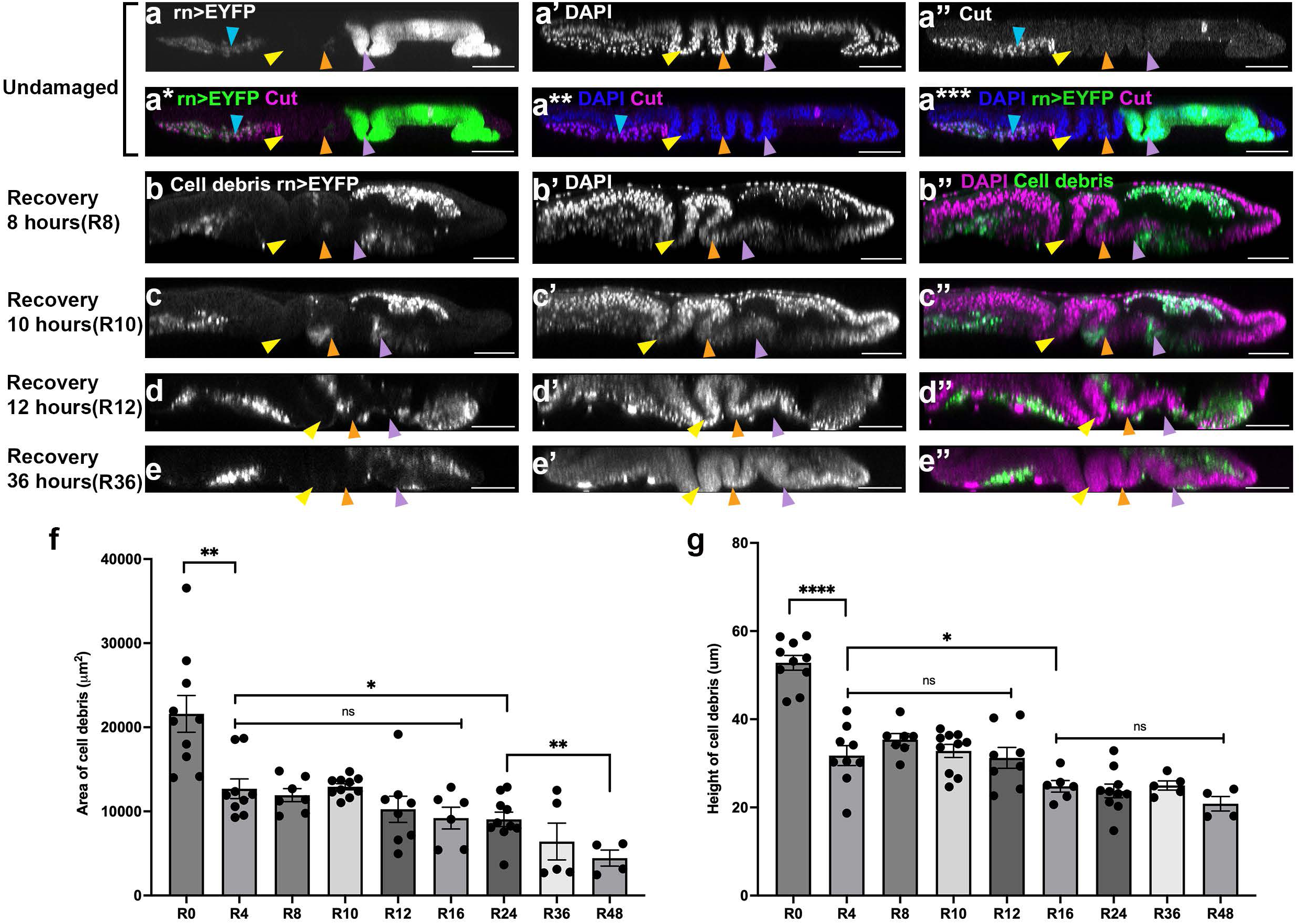
Temporal dynamics of debris clearance in the regenerating wing disc. (a–a***) Orthogonal YZ projections of an undamaged wing disc expressing *rn*>EYFP (a, a*, a***), co-stained with DAPI (a′, a**, a***) and anti-Cut antibody (a″–a***). Light blue arrowheads indicate the adult muscle precursor cells in the notum. (b-e) For all panels, yellow arrowhead: Notum–Hinge fold (N–H); Orange arrowhead: Hinge–Hinge fold (H–H); Purple arrowhead: Hinge–Pouch fold (H–P). (b–e) Orthogonal YZ projections of regenerating discs at R8 (b-b”), R10 (c-c”), R12 (d-d”), and R36 (e-e”), with GFP labeling cell debris (b-e, and b”-e”) and DAPI marking the nuclei (b’-e’, and b”-e”) (f) Quantification of total debris area across regeneration time points. R0 n=10 discs, R4 n=9 discs, **P= 0.0031 compared to R0, R8 n=7 discs, R10 n=10 discs, R12 n=8 discs, R16 n=6 discs, ns P=0.0696 compared to R4, R24 n=10 discs, * P=0.0241 compared to R4, R36 n=5 discs, R48 n=4 discs, ** P=0.0069 compared to R24. (g) Quantification of debris height across regeneration time points, with the same n as in (f). R0-R4 (****P<0.0001), R4-R16 (*P=0.0204). Statistical significance was determined using Welch’s t-test. Scale bars: 50 µm. Error bars: SEM.

**Supplementary Figure S2.**
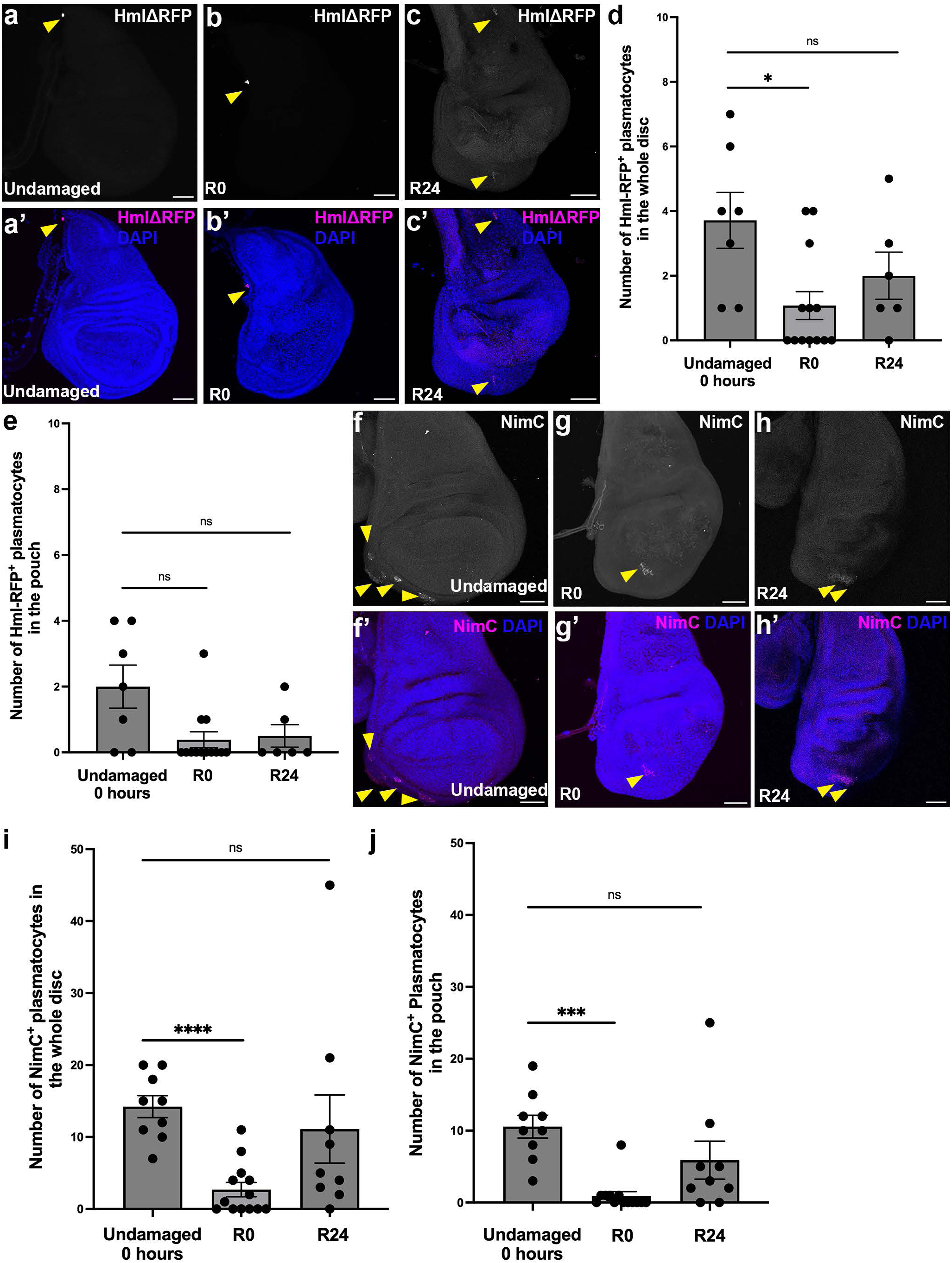
Plasmatocytes are not recruited to the regenerating wing disc epithelium. (a-c′) Wing discs from larvae expressing HmlΔ-RFP co-stained with DAPI. (a-a′) Undamaged control. (b-b′) R0 disc. (c-c′) R24 disc. Yellow arrowheads indicate Hml-RFP-positive plasmatocytes. (d) Quantification of HmlΔ-RFP-positive plasmatocytes throughout the whole wing disc. No damage 0 hours n=7 discs, R0 n=13 discs, *P=0.0231, R24 n=6 discs, ns P=0.1583 (e) Quantification of HmlΔ-RFP-positive plasmatocytes within the wing disc pouch region, same n as in (d). No damage 0 hours-R0, ns P=0.0506. No damage 0 hours-R24, ns P=0.0731. (f-h′) Wing discs co-stained with anti-Nimrod (NimC) and DAPI in undamaged controls (f-f′), at R0 (g-g′), and at R24 (h-h′). Yellow arrowheads indicate NimC-positive plasmatocytes. (i) Quantification of NimC-positive plasmatocytes in the whole wing disc. No damage n=9 discs, R0 n=13 discs, **** P<0.0001, R24 n=9 discs, ns P=0.5459. (j) Quantification of NimC-positive plasmatocytes within the pouch region. No damage n=9 discs compared to R0 n=13 discs, ***P= 0.0002. No damage compared to R24 n=9 discs, ns P=0.1530. Statistical significance was determined using Welch’s t-test. Scale bars: 50 µm. Error bars: SEM.

**Supplementary Figure S3.**
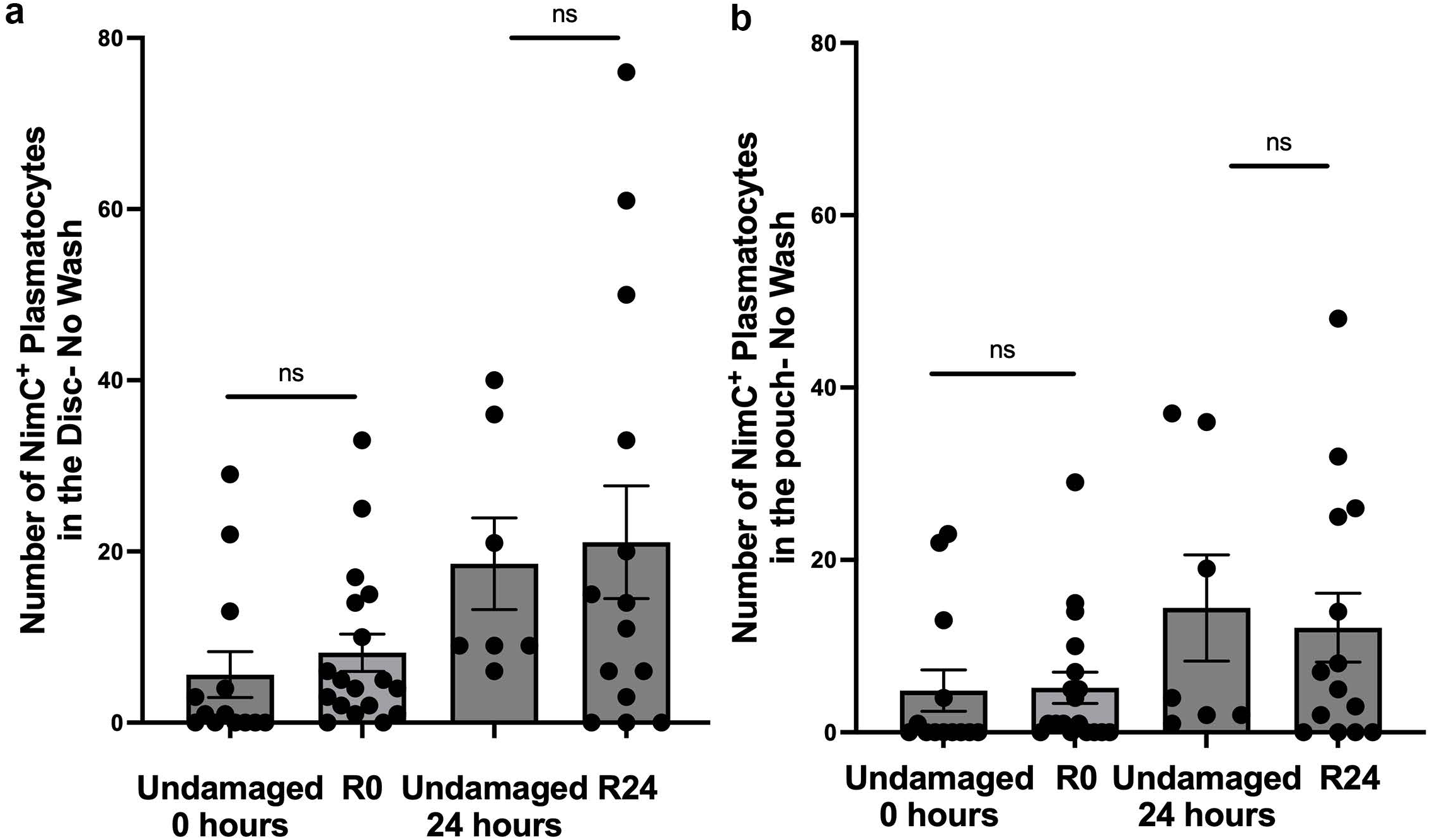
Plasmatocyte quantification under wash and no-wash conditions. (a) Quantification of NimC-positive plasmatocytes in the whole wing disc under no-wash conditions. No damage 0 hours n=13 discs, R0 n=18 discs, ns P= 0.4662. Undamaged 24 hours n=7 discs, and R24 n=14 discs, ns P=0.7715. (b) Quantification of NimC-positive plasmatocytes in wash and no-wash conditions within the wing disc pouch. Wash no damage n= 9 discs, No wash no damage n=13 discs, ns P=0.0610 compared to no damage wash, wash R0 n=13 discs, no wash R0 n=18 discs, * P=0.0373 compared to wash R0, and ns P=0.9158 compared to no damage no wash. Statistical significance was determined using Welch’s t-test. Error bars: SEM.

**Supplementary Figure S4.**
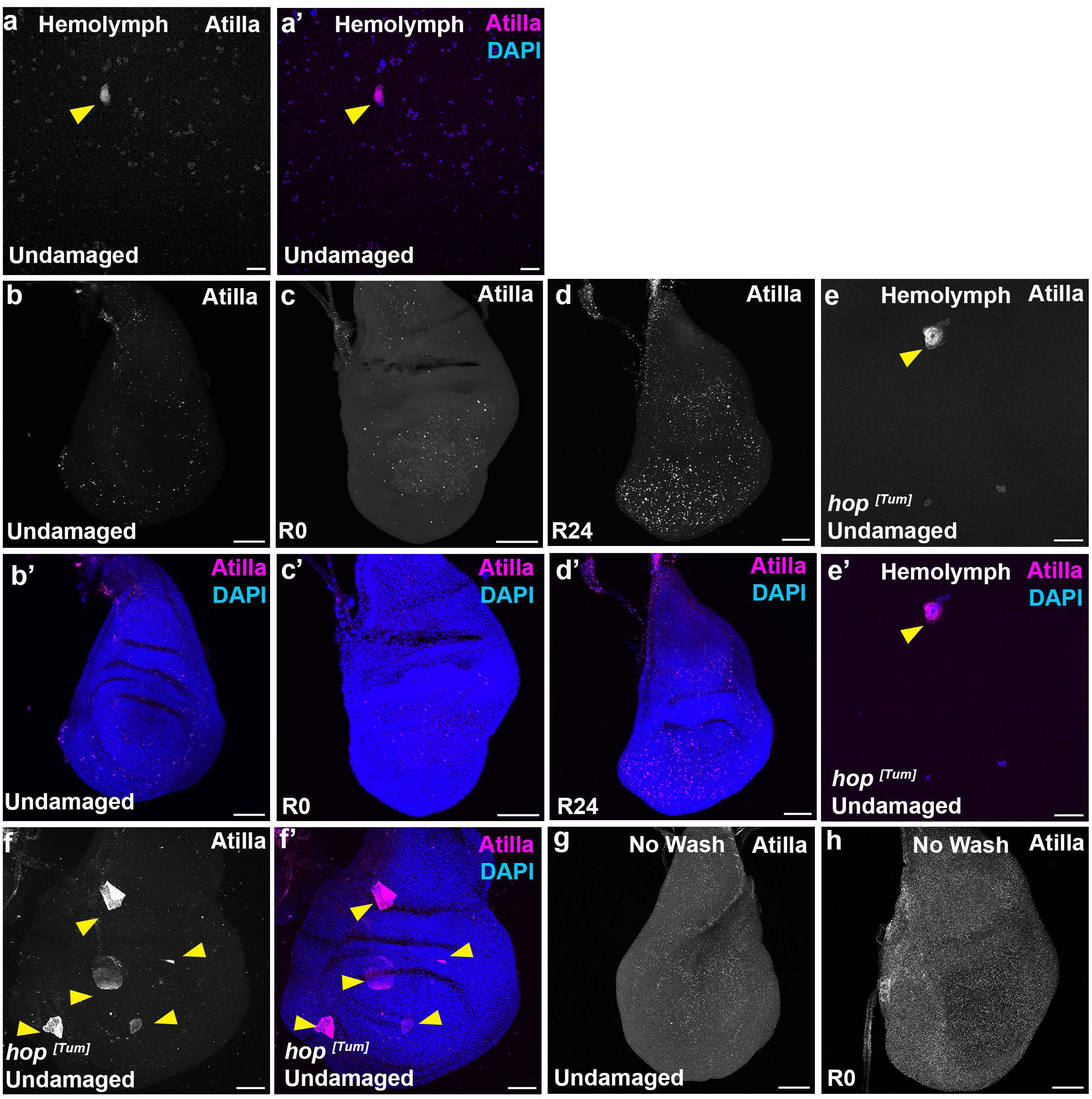
Lamellocytes are not detected in regenerating wing discs. (a–a′) Hemolymph from an undamaged larva containing circulating hemocytes, co-stained with anti-Atilla and DAPI. Yellow arrowheads indicate Atilla-positive lamellocytes. (b–d′) Wing imaginal discs co-stained with anti-Atilla and DAPI. (b–b′) Undamaged controls, 0/15 discs contained lamellocytes. (c–c′) R0, 0/11 discs contained lamellocytes. (d–d′) R24, 0/13 discs contained lamellocytes. (e–e′) Undamaged hop^[Tum]^ larval hemolymph sample co-stained with anti-Atilla and DAPI. The yellow arrowhead indicates an Atilla-positive lamellocyte. (f–f′) Undamaged hop^[Tum]^ wing imaginal disc co-stained with anti-Atilla and DAPI. Yellow arrowheads indicate Atilla-positive lamellocytes (g). Undamaged disc stained with anti-Atilla under no-wash conditions, 0/3 discs contained lamellocytes. (h) R0 disc stained with anti-Atilla under no-wash conditions, 0/3 wing discs contained lamellocytes. Scale bars: 50 µm.

**Supplementary Figure S5.**
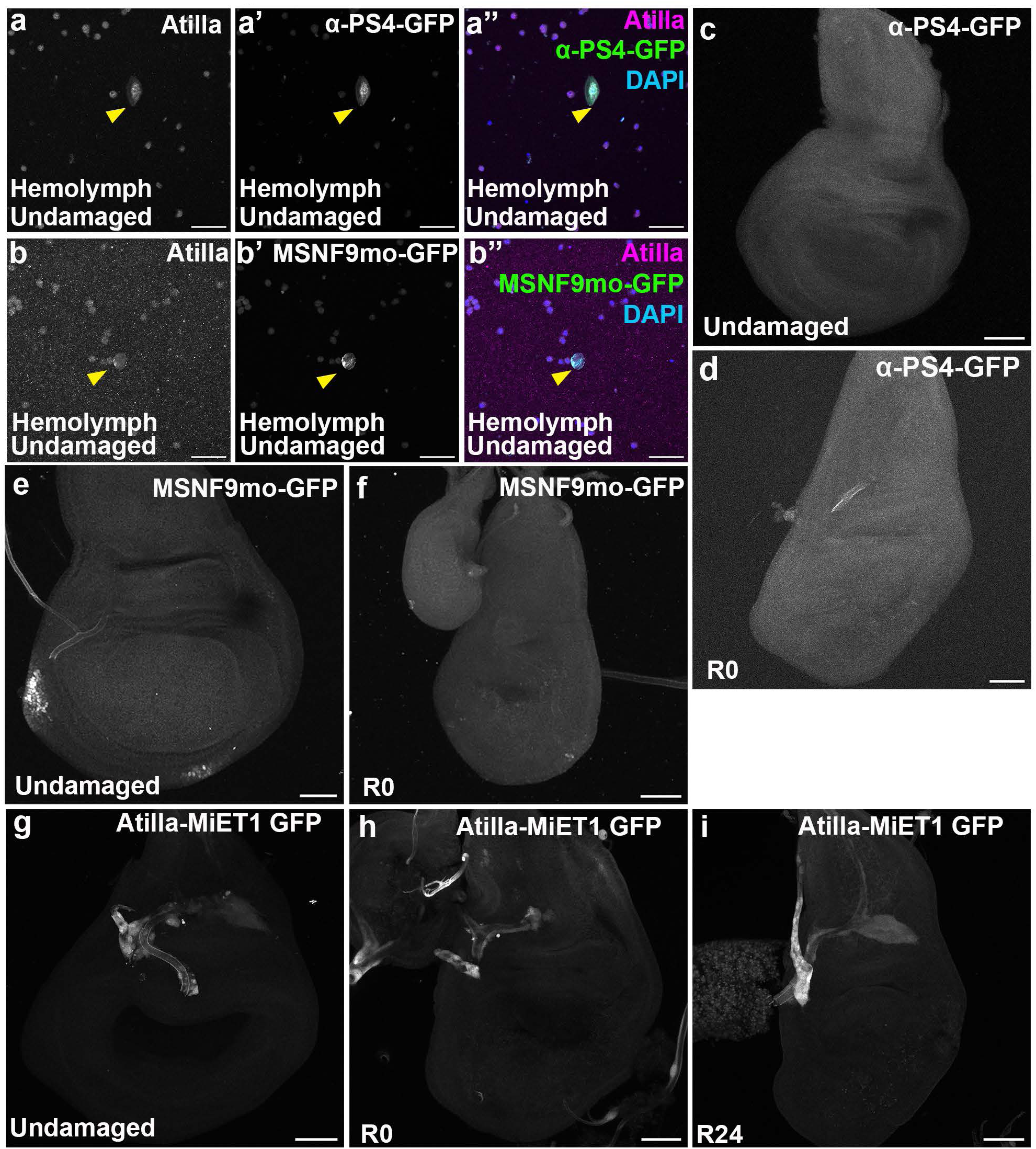
Multiple lamellocyte reporters confirm the absence of lamellocytes in regenerating wing discs. (a-a″) An undamaged larval hemolymph sample containing circulating hemocytes expressing α-PS4-GFP (a′, a″), co-stained with anti-Atilla (a, a″) and DAPI (a″). Yellow arrowheads indicate lamellocytes. (b-b″) Undamaged larval hemolymph sample expressing MSNF9mo-GFP (b′, b″), co-stained with anti-Atilla (b, b″) and DAPI (b″). Yellow arrowheads indicate lamellocytes. (c-d) Top-down images of wing discs from larvae expressing α-PS4-GFP. (c) Undamaged disc, 0/6 discs contained lamellocytes. (d) R0 disc, 0/6 discs contained lamellocytes. (e-f) Top-down images of wing discs from larvae expressing MSNF9mo-GFP. (e) Undamaged disc 0/7discs contained lamellocytes. (f) R0 disc, 0/6 discs contained lamellocytes. (g-i) Top-down images of wing discs from larvae expressing Atilla-MiET1GFP. (g) Undamaged disc, 0/9 discs contained lamellocytes. (h) R0 disc, 0/6 discs contained lamellocytes. (i) R24 disc, 0/5 discs contained lamellocytes. Scale bars: 50 µm.

**Supplementary Figure S6.**
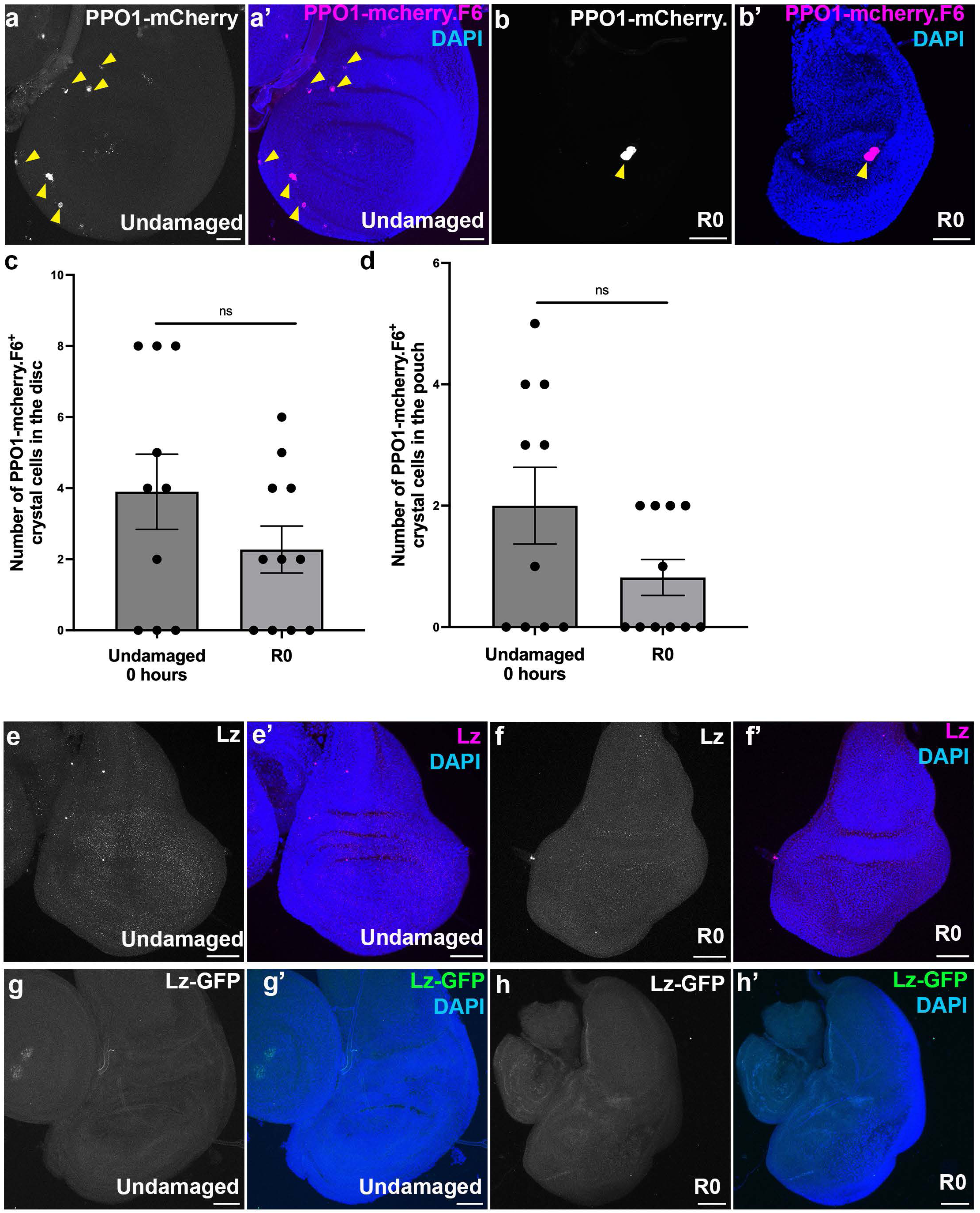
Crystal cells are not enriched in regenerating wing discs. (a-b′) Wing discs from larvae expressing PPO1-mCherry.F6 stained with DAPI. (a-a′) Undamaged control disc (b-b′) R0 disc. Yellow arrowheads indicate PPO1-mCherry.F6-positive crystal cells. (c) Quantification of PPO1-mCherry.F6-positive crystal cells in the whole wing disc. No damage n=10 discs, R0 n=11 discs, ns, P=0.2118 (d) Quantification of PPO1-mCherry.F6-positive crystal cells within the pouch region. No damage-R0 ns, P=0.1147 with the same n as in (c). (e-f′) Top-down images of Wing discs co-stained with anti-Lozenge (Lz) and DAPI in undamaged controls (e-e′) and at R0 (f-f′). (g-h′) Top-down images of wing discs from larvae expressing Lozenge-GFP (Lz-GFP) stained with DAPI. (g-g′) Undamaged control disc. (h-h′) R0 disc. Scale bars: 50 µm. Statistical significance was determined using Welch’s t-test. Error bars: SEM.

**Supplementary Figure S7.**
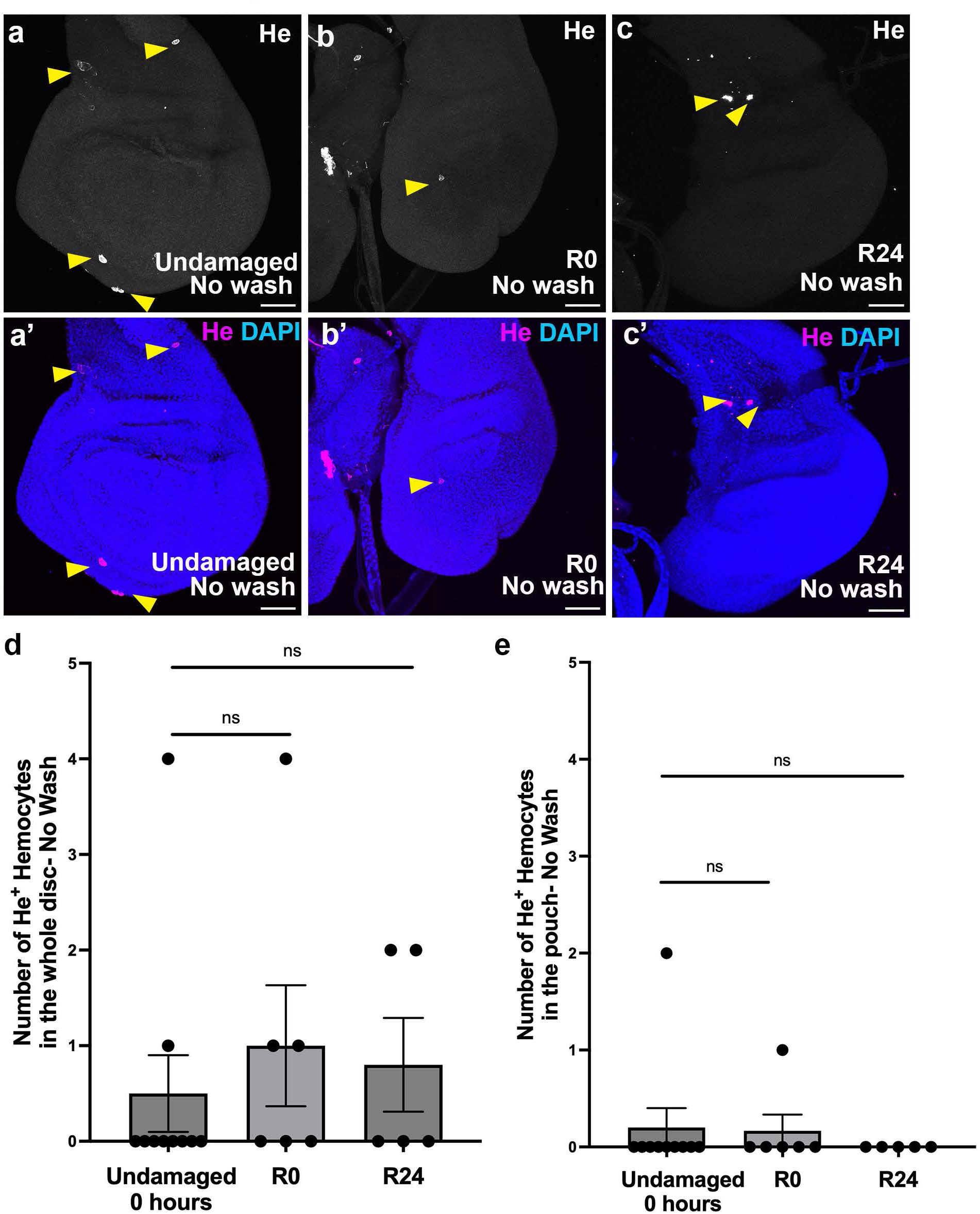
Hemese-positive hemocytes are not enriched in regenerating wing discs. (a-c′) Top-down images of wing discs co-stained with anti-Hemese (He) and DAPI in undamaged controls (a-a′), at R0 (b-b′), and at R24 (c-c′). Yellow arrowheads indicate He-positive hemocytes. (d) Quantification of He-positive hemocytes in the whole wing disc: no damage, n = 10 discs, R0 n = 6 discs, ns, P = 0.5212, and R24 n = 5 discs, ns, P = 0.6466. (e) Quantification of He-positive hemocytes within the pouch region: no damage vs. R0 ns, P = 0.9000, and no damage vs. R24 ns, P = 0.3434, with same n as in (d). Scale bars: 50 µm. Statistical significance was determined using Welch’s t-test. Error bars represent SEM.

**Supplementary Figure S8.**
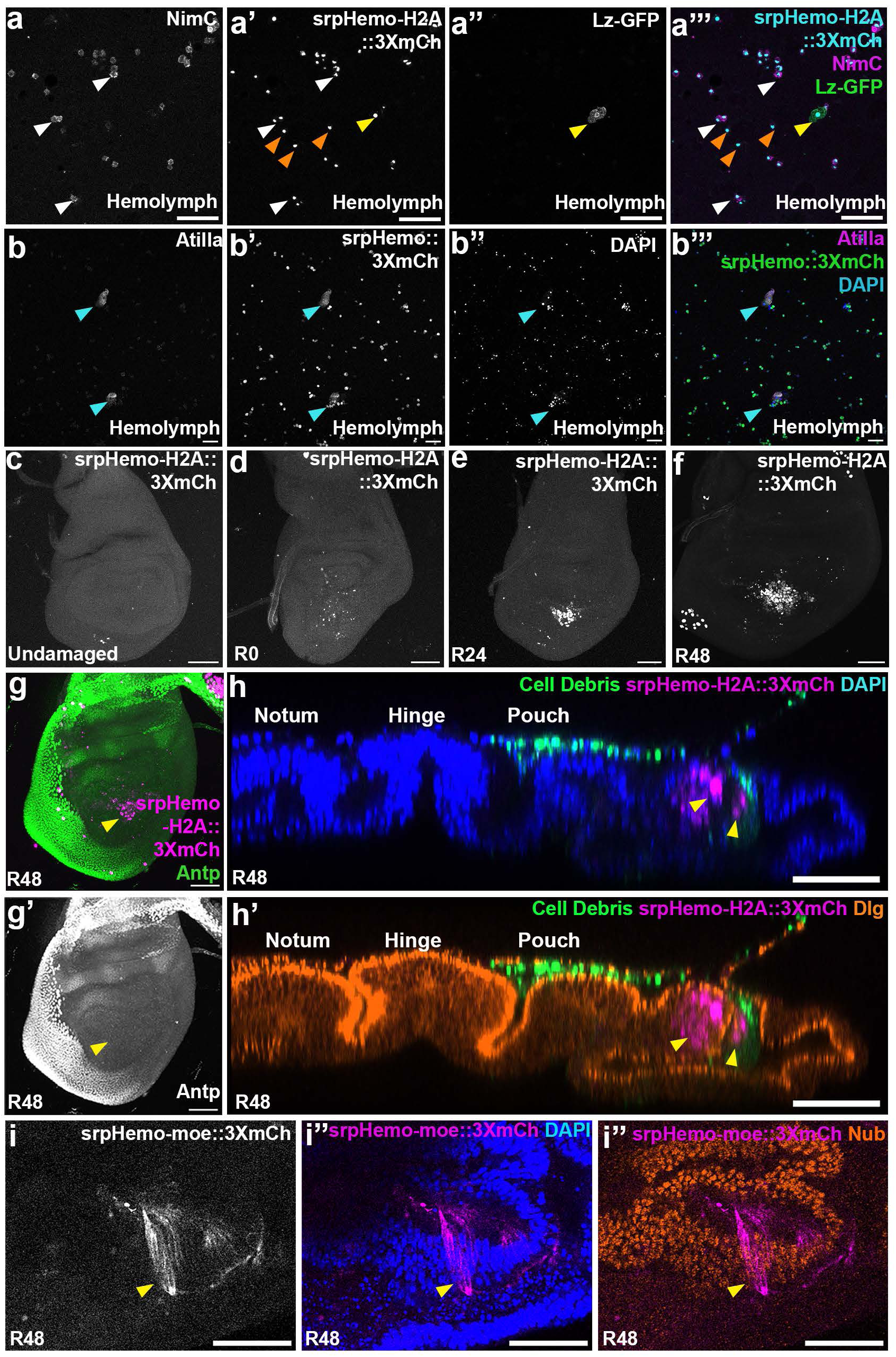
srpHemo reporters reveal columnar epithelial expression during regeneration. (a-a‴) Hemolymph from an undamaged larva containing circulating hemocytes expressing srpHemo-H2A::3XmCh (a′, a‴) and Lz-GFP (a″, a‴), co-stained with anti-Nimrod (a, a‴). The yellow arrowhead indicates a Lz-GFP-positive crystal cell, orange arrowheads indicate srpHemo-H2A::3XmCh-positive hemocytes, and white arrowheads indicate NimC-positive plasmatocytes. (b-b‴) Hemolymph from an undamaged larva expressing srpHemo::3XmCh co-stained with anti-Atilla and DAPI. Blue arrowheads indicate Atilla-positive lamellocytes co-expressing srpHemo::3XmCh. (c-f) Wing discs from larvae expressing srpHemo-H2A::3XmCh. (c) Undamaged control disc. (d) R0 disc. (e) R24 disc. (f) R48 disc. (g-g′) Wing disc from larva expressing srpHemo-H2A::3XmCh at R48 co-stained with anti-Antennapedia (Antp). Yellow arrowhead indicates srpHemo-H2A::3XmCh-positive cells. (h-h′) YZ orthogonal projections of R48 wing discs from larvae expressing srpHemo-H2A::3XmCh to mark putative immune cells and *rn*>EYFP to mark the cell debris co-stained with anti-Discs large (Dlg) to mark cell membranes and DAPI. Yellow arrowheads indicate srpHemo-H2A::3XmCh-positive cells within the regenerating epithelium. (i-i″) R48 wing disc from larvae co-expressing srpHemo-moe::3XmCh to mark membranes of putative immune cells and stained with anti-Nubbin (Nub) to mark the wing pouch and DAPI. Yellow arrowhead labels columnar epithelial pouch cells expressing both the srpHemo-moe::3XmCh reporter and Nubbin. Scale bars: 50 µm.

**Supplementary Figure S9.**
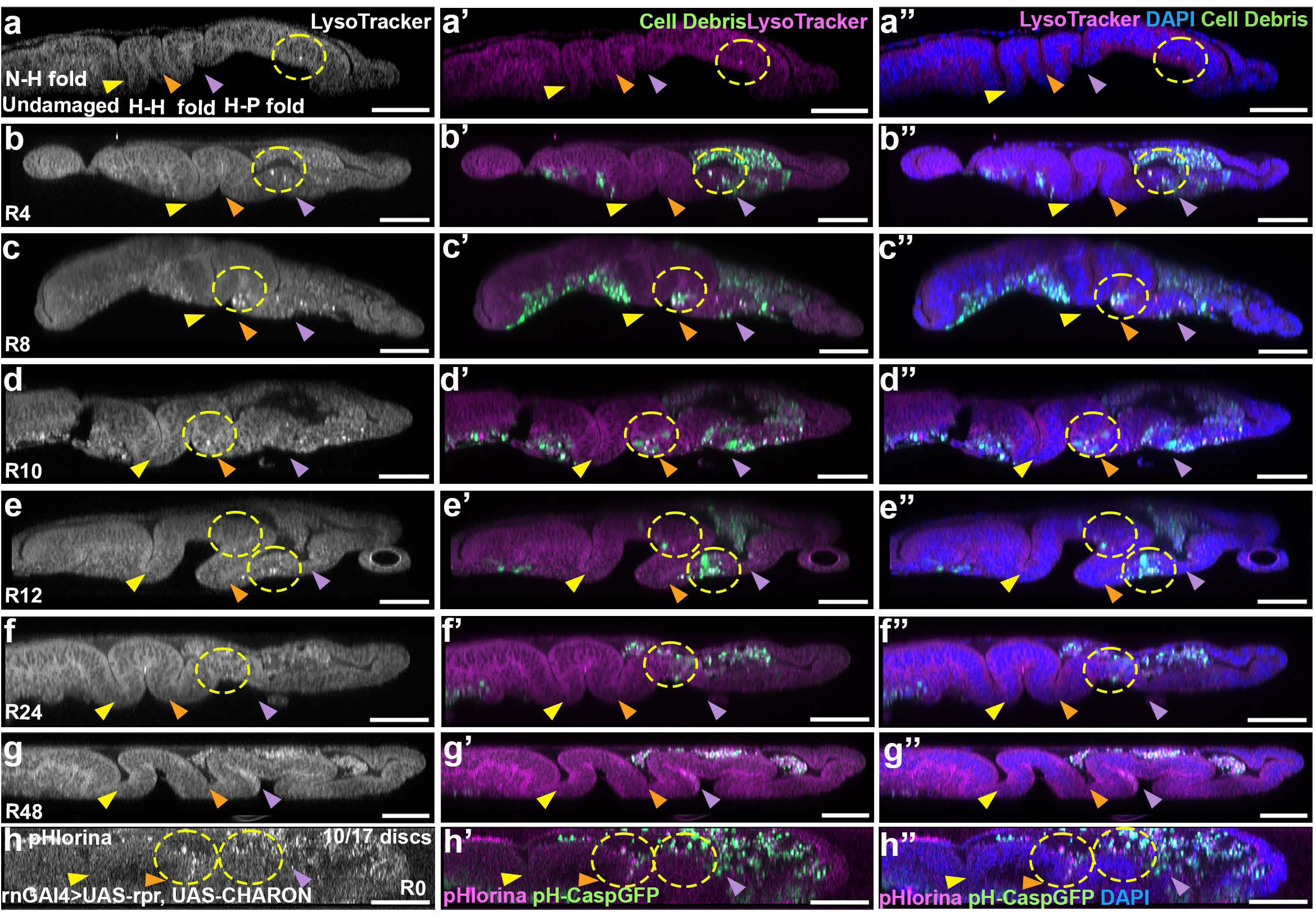
Cell debris clearance is followed by lysosome formation during regeneration. For panels a-f’’, dotted yellow circles mark regions of LysoTracker staining. Yellow arrowhead: Notum-Hinge fold (N-H); Orange arrowhead: Hinge-Hinge fold (H-H); Purple arrowhead: Hinge-Pouch fold (H-P). (a-a″) YZ orthogonal projection of an undamaged wing disc co-stained with LysoTracker (a-a″) and DAPI (a″). (b-b″) YZ orthogonal projection of an R4 wing disc expressing *rn*>EYFP-labeled cell debris (b′, b″), co-stained with LysoTracker (b-b″) and DAPI (b″). (c-c″) YZ orthogonal projection of an R8 wing disc expressing *rn*>EYFP-labeled debris (c′, c″), co-stained with LysoTracker (c-c″) and DAPI (c″). (d-d″) YZ orthogonal projection of an R10 wing disc expressing *rn*>EYFP-labeled debris (d′, d″), co-stained with LysoTracker (d-d″) and DAPI (d″). (e-e″) YZ orthogonal projection of an R12 wing disc expressing *rn*>EYFP-labeled debris (e′, e″), co-stained with LysoTracker (e-e″) and DAPI (e″). (f-f″) YZ orthogonal projection of an R24 wing disc expressing *rn*>EYFP-labeled debris (f′, f″), co-stained with LysoTracker (f-f″) and DAPI (f″). (g-g″) YZ orthogonal projection of an R48 wing disc expressing *rn*>EYFP-labeled debris (g′, g″), co-stained with LysoTracker (g-g″) and DAPI (g″). (h-h″) *rn*GAL4>UAS-rpr, UAS-CHARON-expressing wing imaginal disc at R0 stained with DAPI (h″), showing pHlorina (h-h″) and pH-CaspGFP (h′, h″) signals. 10/17 discs were positive for phlorina signal in the hinge and pouch region. Dotted yellow circles mark regions of pHlorina expression. Scale bars: 50 µm.

**Supplementary Figure S10.**
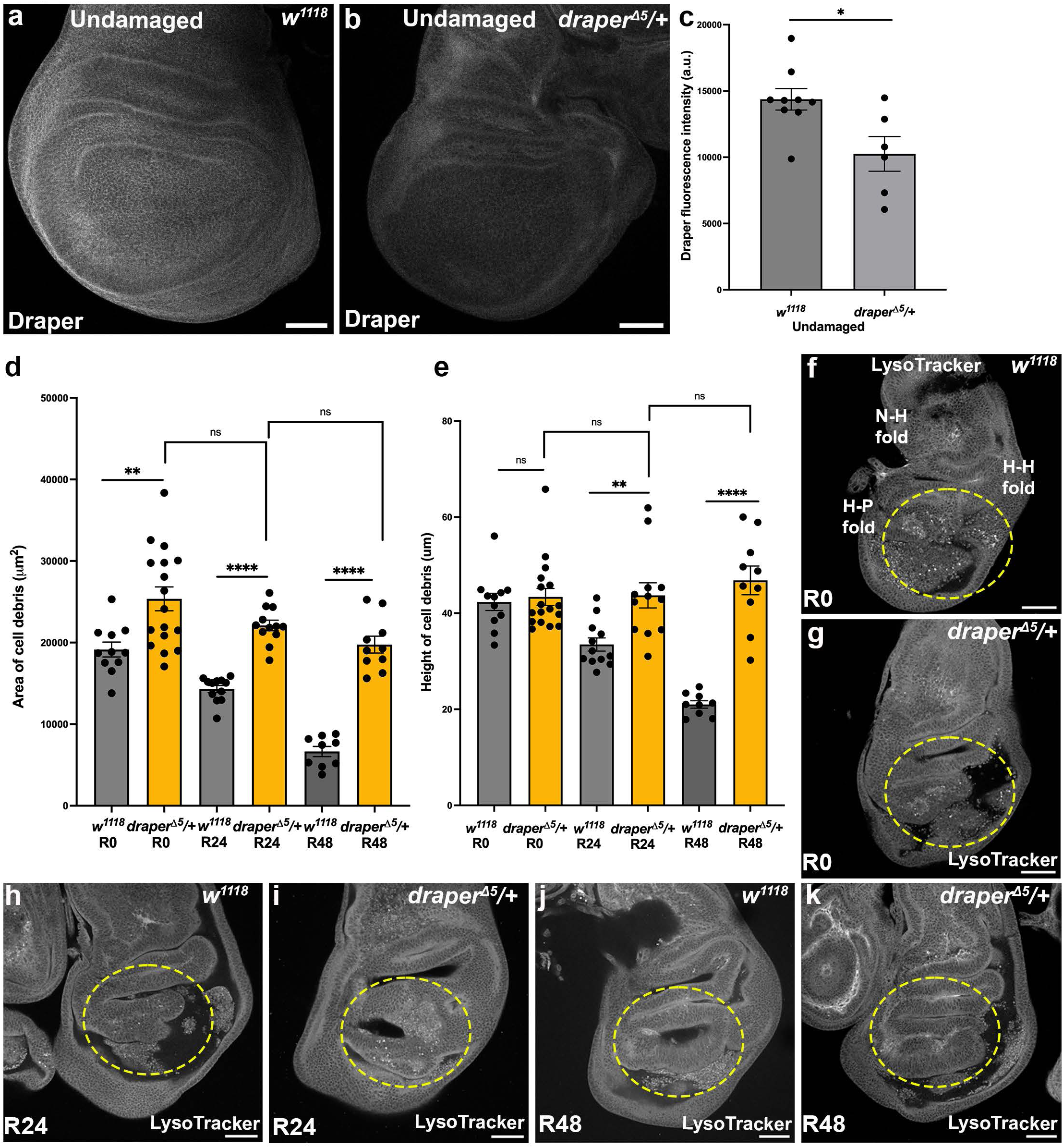
Heterozygous *draper* mutants show increased debris retention and delayed lysosome dynamics during regeneration. (a-b) Undamaged *w^1118^* (a) and *draper^Δ5^*/+ (b) wing discs stained with anti-Draper. (c) Quantification of total Draper fluorescence intensity in *w^1118^* and *draper^Δ5^*/+ wing discs. *w^1118^* n=9, and *draper^Δ5^*/+ n=6, *P=0.0259. (d) Quantification of total debris area in *w^1118^* (grey bars) and *draper^Δ5^*/+ (yellow bars) wing discs across regeneration time points. *w^1118^* R0 n=11 discs, *draper^Δ5^*/+ R0 n=17 discs, **P=0.0014, *w^1118^* R24 n=12 discs, *draper^Δ5^*/+ R24 n=12 discs,****P<0.0001, *draper^Δ5^*/+ R0 and *draper^Δ5^*/+ R24 ns, P=0.0537, *w^1118^* R48 n=9 discs, *draper^Δ5^*/+ R48 n=10 discs, ****P<0.0001, and *draper^Δ5^*/+ R24 and *draper^Δ5^*/+ R48 ns, P=0.0718. (e) Quantification of debris height in *w^1118^* (grey bars) and *draper^Δ5^/+* (yellow bars) wing discs across regeneration time points. *w^1118^* R0 n=11, and *draper^Δ5^*/+ R0 n=17 discs, ns, P=0.6836, *w^1118^* R24 n=12 discs, and *draper^Δ5^*/+ R24 n=12 discs, **P=0.0031, *w^1118^* R48 n=9 discs, and *draper^Δ5^*/+ R48 n=10 discs, ****P<0.0001. Pairwise comparisons were not significant between draperΔ5/+ R0 and draperΔ5/+ R24, ns, P=0.9222, and draperΔ5/+ R24 compared to R48, ns, P=0.4409. (f-k) *w^1118^* and *draper^Δ5^/+* wing discs stained with LysoTracker at R0 (f, g), R24 (h, i), and R48 (j, k). Yellow dotted circles indicate lysosome-positive regions. Notum-Hinge fold (N-H), Hinge-Hinge fold (H-H), and Hinge-Pouch fold (H-P) are marked in (f). Scale bars: 50 µm. Statistical significance was determined using Welch’s t-test. Error bars represent SEM.

**Supplementary Figure S11.**
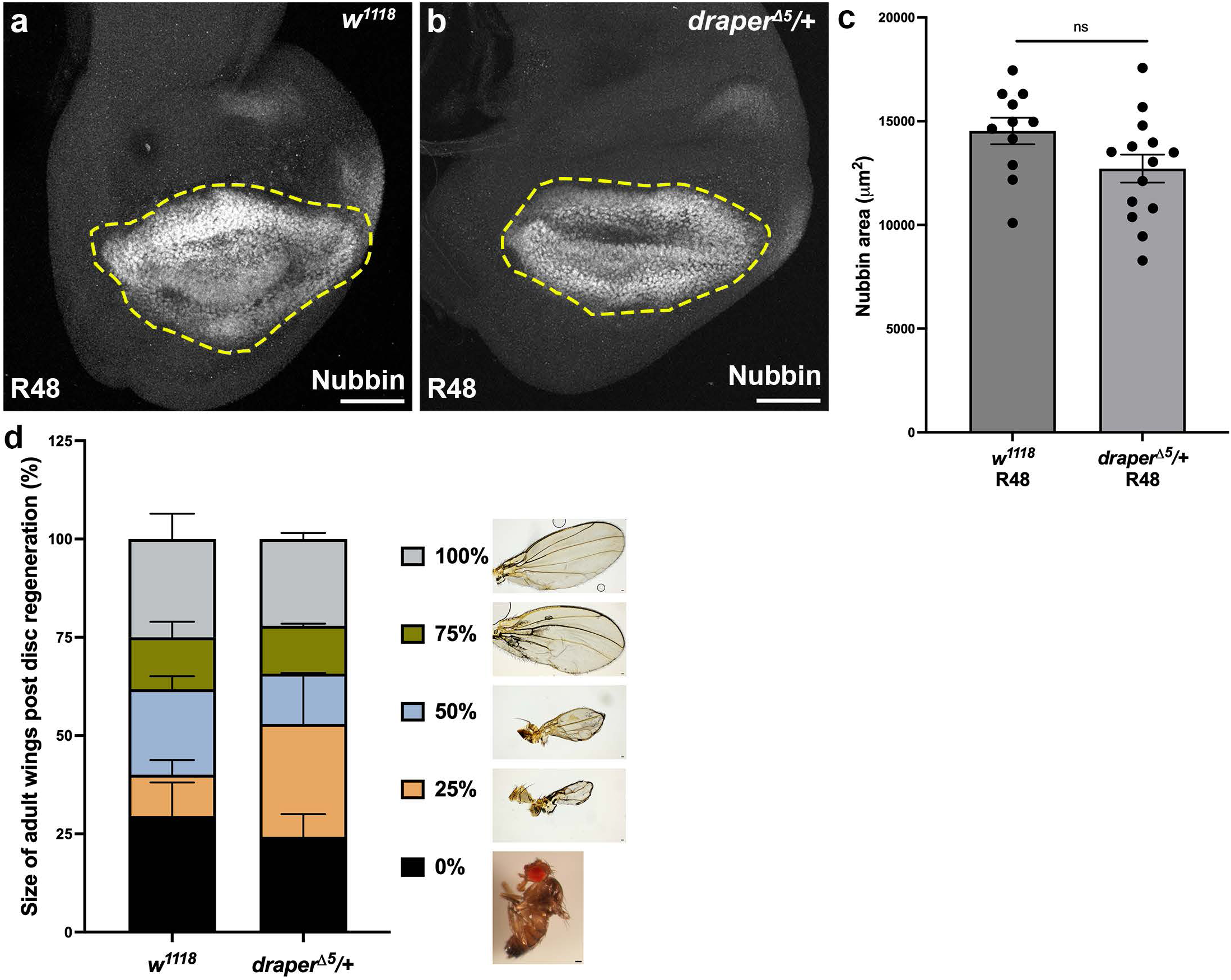
Heterozygous *draper* mutants impair overall regeneration. (a-c) R48 wing discs stained with anti-Nubbin to mark the regenerating pouch in *w^1118^* (a) and *draper^Δ5^*/+ (b) discs. (c) Quantification of total Nubbin-positive pouch area in R48 *w^1118^* wing discs, n=11 discs, and *draper^Δ5^*/+ wing discs, n=14 discs, ns, P=0.0642). Dotted yellow circles indicate the Nubbin area. Statistical significance was determined using Welch’s t-test. (d) Quantification of adult wing size following disc regeneration in *w^1118^* and *draper^Δ5^*/+ mutants. *w^1118^* n=302 wings, *draper^Δ5^*/+ n=986 wings, *P=0.0142. Statistical significance was determined using chi-square test. Scale bars: 50 µm. Error bars represent SEM.

